# cDC1 and interferons promote spontaneous CD4^+^ and CD8^+^ T cell protective responses to breast cancer

**DOI:** 10.1101/2020.12.23.424092

**Authors:** Raphaël Mattiuz, Carine Brousse, Marc Ambrosini, Jean-Charles Cancel, Gilles Bessou, Julie Mussard, Amélien Sanlaville, Christophe Caux, Nathalie Bendriss-Vermare, Jenny Valladeau-Guilemond, Marc Dalod, Karine Crozat

## Abstract

Here we show that efficient breast cancer immunosurveillance relies on cDC1, conventional CD4^+^ T cells, CD8^+^ cytotoxic T lymphocytes (CTL) and later NK/NK T cells. For this process, cDC1 were required constitutively, but especially during the T cell priming phase. In the tumor microenvironment, cDC1 interacted physically and jointly with both CD4^+^ T cells and tumorspecific CD8^+^ T cells. We found that interferon (IFN) responses were necessary for the rejection of breast cancer, including cDC1-intrinsic signaling by IFN-γ and STAT1. Surprisingly, cell-intrinsic IFN-I signaling in cDC1 was not required. cDC1 and IFNs shaped the tumor immune landscape, notably by promoting CD4^+^ and CD8^+^ T cell infiltration, terminal differentiation and effector functions. XCR1, CXCL9, IL-12 and IL-15 were individually dispensable for breast cancer immunosurveillance. Consistent with our experimental results in mice, high expression in the tumor microenvironment of genes specific to cDC1, CTL, helper T cells or interferon responses are associated with a better prognosis in human breast cancer patients. Our results show that immune control of breast cancer depends on cDC1 and IFNs as previously reported for immunogenic melanoma or fibrosarcoma tumor models, but that the underlying mechanism differ. Revisiting cDC1 functions in the context of spontaneous immunity to cancer should help defining new ways to mobilize cDC1 functions to improve already existing immunotherapies for the benefits of patients.

**Synopsis:** Type 1 conventional dendritic cells cross-present tumor antigens to CD8^+^ T cells. Understanding the regulation of their antitumor functions is important. Cell-intrinsic STAT1/IFN-γ signaling licenses them for efficient CD4^+^ and CD8^+^ T cell activation during breast cancer immunosurveillance.

## Introduction

Conventional dendritic cells (cDC) are specialized in antigen (Ag) capture, processing and presentation for T cell priming (1). cDC are present in lymphoid organs and peripheral tissues. Lymphoid resident-cDC (Res-cDC) of the spleen and lymph nodes (LNs) participate in the capture of Ag from blood and lymph respectively. cDC in non-lymphoid tissues capture Ag at the periphery and migrate to the LNs via the afferent lymphatic vessels. cDC encompass two distinct cell types. Type 1 cDC (cDC1) excel in cytotoxic CD8^+^ T cell (CTL) activation, particularly via Ag cross presentation (2–4). Type 2 cDC (cDC2) are particularly effective for helper CD4^+^ T cell activation (1).

Previous studies suggested that cDC1 play a non-redundant role in anti-tumor immunity, both for spontaneous control of syngeneic tumor grafts used as a surrogate model for cancer immunosurveillance, and for rejection of established tumors upon immunotherapy (5,6). Yet, most of these studies used mutant mice whose deficiencies were not affecting exclusively cDC1. *Irf8* deficiency in CD11c-expressing cells also affected the differentiation and functions of plasmacytoid dendritic cells (7) and inflammatory cDC2 (8). Batf3 knock-out does not only abrogate cDC1 differentiation (9) but was also recently shown to enhance CD4^+^ regulatory T cell (Treg) induction (10) and to hamper CTL survival and memory (11,12) in a cell-intrinsic manner. Hence, mouse models targeting cDC1 with a higher specificity are mandatory to investigate whether and how they contribute to antitumor immunity (11). This is the case of mice knocked-in for the Cre recombinase in the *Xcr1* locus (13).

A number of key features of cDC1 are proposed to contribute to their critical role in antitumor immunity, beyond their efficiency at cross-presenting cell-associated antigens (5,6). cDC1 may have the unique ability to simultaneously deliver to CTLs a series of complementary output signals ensuring their optimal response. CXCL9 attracts CXCR3-expressing memory or effector CTLs to the tumors. IL-12 production and IL-15 trans-presentation promote CTL IFN-γ expression and proliferation. cDC1 could also promote delivery to CTLs of help from other immune cells including CD4^+^ T cells (5,14). However, which of these output signals are critical for cDC1 antitumor immunity remains to be rigorously investigated (5,15).

The antitumor functions of cDC1 has been proposed to depend on their integration of specific input signals, instructing them to deliver the right output signals to CTLs. cDC1 recruitment into the tumor bed can be promoted by engagement of their chemokine receptor Xcr1 (16), whose ligand Xcl1 is produced by activated NK cells and CTLs (17). Triggering of the receptor for type I interferons (IFNAR) on cDC1 is necessary for rejection of immunogenic melanoma and fibrosarcoma by promoting Ag cross-presentation (18,19). It also boosts their expression of co-stimulation molecules, induces their trans-presentation of IL-15, and can promote their production of IL-12 (5,20). However, whether these input signals are always required by cDC1s for their antitumor functions is unknown. Moreover, to which extent and how different types of interferons promote the immunogenic maturation of cDC1, not only IFN-I but also IFN-III or IFN-γ, remains to be formally investigated.

Here, we studied whether cDC1 promote spontaneous immunity to breast cancer, and when, where and how this is achieved. To this aim, we used a mouse model of spontaneous immune control of breast cancer in C57BL/6 female mice, consisting of an orthotopic graft of a subclone of the NOP23 syngeneic breast adenocarcinoma cell line expressing epitopes from the ovalbumin (OVA) model antigen (21). This experimental system enabled us to study the cellular and molecular mechanisms underpinning spontaneous immune control of breast cancer, by harnessing the *Karma-tmt-hDTR* (22), *Xcr1-DTA, Xcr1^Cre^* and *Karma^Cre^* (13) mutant mouse models that we have generated and validated to specifically target cDC1, in combination with other mutant C57BL/6 mice.

## Materials and Methods

### Ethics statement regarding care and use of animals for experimentation

Mice were bred and maintained the CIPHE pathogen-free animal facility. The study was carried out in accordance with institutional guidelines and with protocols approved by the Comité National de Réflexion Ethique sur l’Expérimentation Animale #14 and the Ministère de l’Enseignement Supérieur, de la Recherche et de l’Innovation (ROXinAIR APAFiS #1221 and #16555).

### Mice and in vivo treatments

All experiments were performed with female littermate mice at 7–15 weeks of age. The mouse strains used are all on the C57BL/6J background and listed in Table S1. For a sustained and efficient cDC1 conditional depletion for at least 10 consecutive days, *Karma-tmt-hDTR* and *Xcr1^Cre/wt^;Rosa26^hDTR/wt^* mice received a first dose of 32 ng/g of body weight of DT (Merck), followed by one injection of 16 ng/g every 60h. For *in vivo* antibody-mediated cell depletion, C57BL/6 mice were injected i.p. with the antibody and doses indicated in Table S2, starting 1d before tumor engraftment and then as indicated in the figures. To block lymphoid cell egress from peripheral lymphoid organs, mice received 20 μg of FTY720 (Cayman Chemical) starting 1d before tumor engraftment, then every 2d.

### Tumor experiments

We used a breast adenocarcinoma tumor cell line that is spontaneously rejected when orthotopically implanted in C57BL/6 females. It was derived from the NOP23 cell line, which was established from a spontaneous mammary tumor of a transgenic mouse expressing a dominant negative version of p53 and the rat NEU (HER2) oncogene fused at its COOH terminus to class I and II OVA peptide sequences (21). NOP23 cells were grown in DMEM, 10% FCS, supplemented with 10mg/ml of Insulin transferrin Sodium Selenite media supplement (Sigma-Aldrich). 5.10^6^ NOP23 cells were injected in the mammary inguinal fat pad, under isoflurane anesthesia. Tumor volume was calculated as 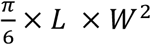. where *L* is the greatest length and *W* is the width of the tumor, measured with a caliper. Graphs of tumor volume are represented as mean+/−SEM.

### Shield bone marrow chimera mice

To generate *Ifngr1^−/−^→Xcr1*-DTA and *Stat1^−/−^→Xcr1-DTA* shield bone marrow chimera (SBMC) mice, the hind legs of *Xcr1*-DTA recipient mice were irradiated with one dose of 8 Gy, to preserve most of their hematopoietic system that remained WT. The cDC1 empty niche of these recipient mice was then reconstituted upon injection of 30.10^6^ bone marrow cells from *Ifngr1^−/−^* or *Stat1^−/−^* donor mice. *Xcr1*-DTA→*Xcr1*-DTA and WT→*Xcr1*-DTA SBMC mice were used as controls. NOP23 cells were engrafted 8 to 14 weeks later.

### Preparation of cell suspension from tumors and TdLNs for flow cytometry

Tumors and tumor-draining inguinal LNs (TdLNs) were cut into small pieces with a scalpel and incubated in a Collagenase D (1 mg/ml)/ of DNase I (70 μg/ml) enzymatic cocktail (Roche) in RPMI, for 30min at 37°C. Ice-cold PBS/EDTA (2 mM) was added for 5 min. Digested tissues were crushed through a 70 μm nylon sieve. For flow cytometry, cells were pre-incubated with 2.4G2 mAb to block Fc-receptors, then stained with the mAb listed in Table S2 for 25 min at 4°C. Class-I OVA Tetramer (iTAg Tetramer/PE-H-2 Kb OVA [SIINFEKL], MBL International) was incubated at 1/100 at 4°C for 1 hour, and Class-II OVA Tetramer (T-Select I-Ab OVA323-339 [ISQAVHAAHAEINEAGR] Tetramer-APC, MBL International) at 1/12.5 at room temperature for 1 hour, before proceeding to staining with mAbs. For CCR7 staining, cells were incubated for 30 min at 37°C. For intracellular staining, cells were re-stimulated *ex vivo* for 4h at 37°C with 0.5 μg/ml of SIINFEKL peptide and 10 μg/ml of Brefeldin A (Sigma-Aldrich) in complete RPMI. Cytokines, Ki67 and FoxP3 were stained after fixation/permeabilization with FoxP3/Transcription Factor Staining Buffer Set (eBioscience). Data were acquired on a LSRFortessa X-20 flow cytometer (BD Biosciences), and analyzed using FlowJo (Tree Star, Inc.). Flow cytometry heatmaps were performed using the Morpheus website from the Broad Institute (https://software.broadinstitute.org/morpheus/).

### Immunohistofluorescence

1,000 naive GFP-expressing OT-I cells were purified from a *Tg^TcraTcrb1100Mjb^;Rag2^−/−^;Ubc-GFP^+/+^* spleen with the Dynabeads Untouched Mouse CD8 Cells Kit (ThermoFisher Scientific) and transferred i.v. in *Xcr1^Cre/wt^;Rosa26^tdRFP/wt^* mice 1d before tumor engraftment. 7d after, tumors were harvested and 12-μm frozen sections were stained with the antibodies listed in Table S2, as described previously (23).

### Quantitative PCR analysis

Total RNA from tumors and TdLNs were prepared with the RNeasy Plus mini kit (QIAGEN). RNA was reverse transcribed into cDNA using the QuantiTect reverse transcription kit (QIAGEN). qPCR was performed with the SybrGreen kit (Takara) and specific primers (Table S3), and run on a 7500 Real Time PCR System apparatus (Applied Biosystems). Relative gene expression was calculated using the ΔΔCt method with *Hprt* as housekeeping control gene.

### Transcriptomic data from breast cancer patients

*XCR1* Kaplan Meier plot was obtained from the Kaplan Meier-plotter database (https://kmplot.com/analysis/), and *CCR7* and *CCL19* Kaplan Meier plots from The Cancer Genome Atlas (TCGA) database. Transcripts enriched in tumors of breast cancer patients with a better overall survival (367 genes) or with a poor prognosis (210 genes) from TCGA were extracted through the human protein atlas (https://www.proteinatlas.org/humanproteome/pathology). A gene is considered prognostic if correlation analysis of gene expression and clinical outcome resulted in Kaplan-Meier plots with high significance (p<0.001). The gene ontology analyses on the good and bad prognosis gene lists were performed with DAVID 6.8 (https://david.ncifcrf.gov/).

### Statistical analyses

Statistical analyses were performed using unpaired Student’s *t*-tests or nonparametric Mann-Whitney tests (MW) when specified. N.S., non-significant (P>0.05); *, P≤0.05; **, P≤0.01; ***, P≤0.001, ****, P≤0.0001.

## Results

### CTL, CD4^+^ T_conv_ and NK1.1^+^ cells are instrumental to rejection of NOP23 mammary tumors

We first investigated whether T or NK cells were providing the effector arm of the spontaneous rejection of the NOP23 breast adenocarcinoma cells in C57BL/6J females. Tumor rejection was abolished by continuous depletion of CTLs or of NK1.1^+^ cells *(i.e.* NK cells and a fraction of NKT cells) (**Fig. 1A**). However, the anti-NK1.1 mAb-mediated depletion showed a delayed and milder effect than CTL depletion. Continuous depletion of all CD4^+^ T cells also abrogated tumor control (**Fig. 1B**), whereas this was not the case of the selective depletion of intra-tumor Treg as achieved with administration of the anti-CTLA-4 clone 9D9 (24). Thus, CTLs, CD4^+^ conventional T cells (T_conv_) and NK/NK T cells are required for NOP23 tumor rejection.

**Figure 1.**
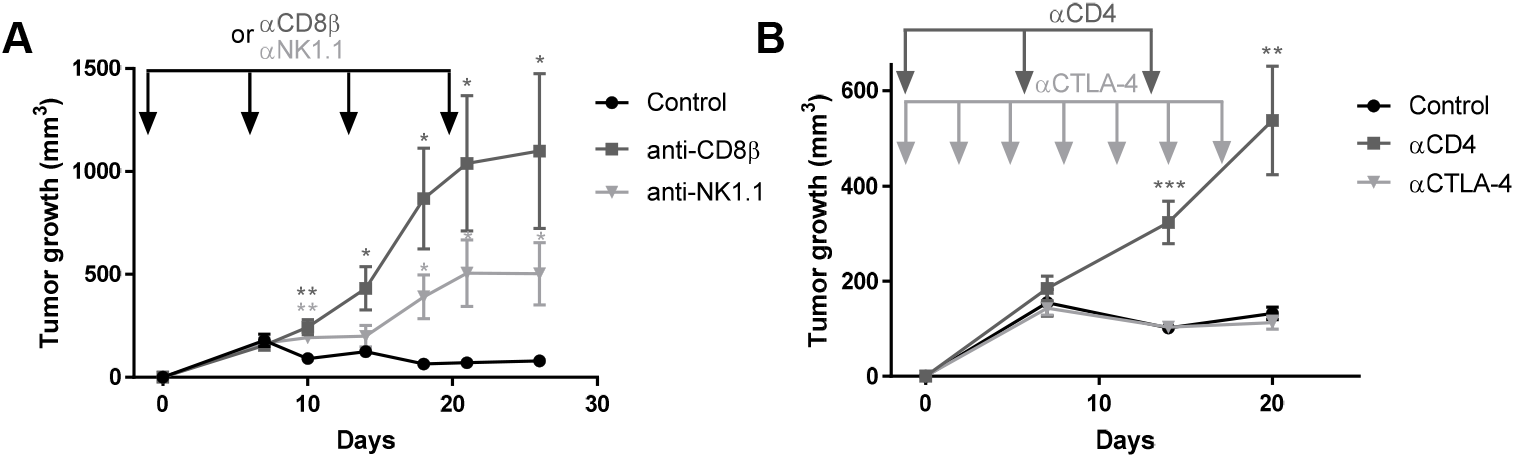
CTL, CD4^+^ T_conv_ and NK1.1^+^ cells are instrumental in spontaneous rejection of breast cancer NOP23. **A**, NOP23 tumor growth in the mammary fat pad of female mice treated or not with anti-CD8β or anti-NK1.1 mAb at the indicated time (arrows) with the first injection given 1d before tumor engraftment. One experiment representative of at least 2 independent ones with 5 mice per group is shown. **B**, NOP23 tumor growth in female mice treated or not with anti-CD4 (once a week) or anti-CTLA-4 (every 3 days) mAb at the indicated time (arrows) with the first injection given 1d before tumor engraftment. n=7 mice per group. *, p < 0.05; **, p < 0.01; ***, p < 0.001 (unpaired *t* test).

### cDC1 are critical for breast cancer control, especially during the T cell priming phase

We next examined, whether and when cDC1 depletion compromised the immune control of NOP23 growth, by taking advantage of our mutant mouse models specifically targeting cDC1. The *Xcr1^Cre/wt^;Rosa26^DTA/wt^ (Xcr1-DTA)* model (Mattiuz et al., 2018) harbored a constitutive and complete lack of all cDC1 in the spleen and IngLNs (**Fig. S1A-B**). The *Xcr1^Cre/wt^;Rosa26^hDTR/wt^ (Xcr1-hDTR)* mouse model allowed conditional depletion of cDC1 in spleen, and of Mig-cDC1 and Res-cDC1 in the inguinal lymph nodes (IngLNs), for at least 2 days following the administration of a single dose of diphtheria toxin (DT) (**Fig. S1A-B**), similarly to the *Karma-tmt-hDTR (Karma)* mice (Alexandre et al., 2016). None of the other immune cell types examined in the spleen and IngLNs were affected in these mice (**Fig. S1C**). *Xcr1-DTA* mice harbored a progressive and unabated growth of NOP23 as compared to control WT animals (**Fig. 2A**), demonstrating that cDC1 are necessary to efficiently reject this breast adenocarcinoma. To determine when cDC1 were required to promote this rejection, we conditionally depleted these cells in *Karma* mice during different time windows relative to tumor engraftment (**Fig. 2A**). The earlier the depletions were initiated, the stronger the tumors grew. However, the tumors never grew as strongly as in the *Xcr1-DTA* mice that are constitutively devoid of cDC1 (**Fig. 2A-B**). TdLN mig-cDC1 showed a high expression of the co-stimulatory molecules CD40 and CD86 at day 4 post-engraftment, as compared to Res-cDC1 or to Mig-cDC1 on any other days (**Fig. 2C**). In summary, cDC1 were required continuously until tumor rejection, but their presence was especially critical during the first 4 days after tumor engraftment, most likely during the phase of T cell priming.

**Figure 2.**
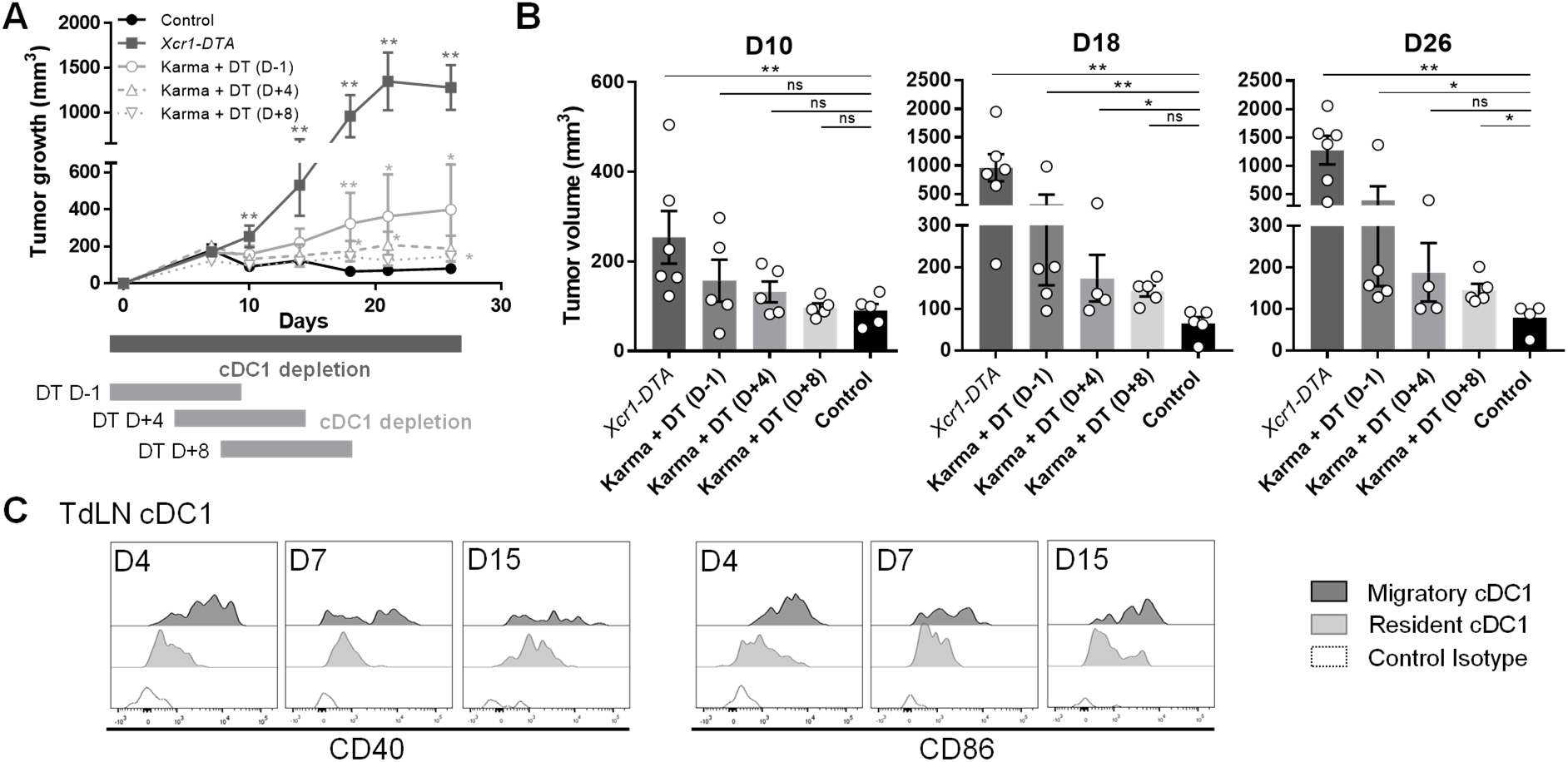
cDC1 are instrumental in spontaneous rejection of breast cancer NOP23, especially during the phase of T cell priming. **A**, Tumor growth in control (n=5), in constitutively cDC1-depleted *(Xcr1-DTA,* n=6) or conditionally cDC1-depleted *(Karma-tmt-DTR* + DT) female mice. *Karma-tmt-DTR* mice were injected 4 times with DT every 60h, starting 1d before engraftment (n=5, representative of 3 independent datasets), at d+4 post-engraftment (n=5, representative of 2 independent datasets), or at d+8 post engraftment (n=5, representative of 2 independent datasets). **B**, Tumor volumes as measured in panel (A) at day 10, 18 and 26. ns, not significant (p > 0.05); *, p < 0.05; **, p < 0.01; ***, p < 0.001; (non-parametric Mann-Whitney test). **C,** Analysis of CD40 and CD86 expression by flow cytometry on TdLN Mig-cDC1 and Res-cDC1. The data shown are from one experiment representative of two independent ones.

### CCR7- and S1PR1-dependent immune cell trafficking is instrumental to NOP23 rejection

At day 7 post-engraftment, CCR7 was upregulated on cDC1 that had migrated from the periphery into the TdLN (**Fig. S2A**). We thus investigated whether CCR7-dependent immune cell trafficking was critical for NOP23 rejection. In *Ccr7^−/−^* mice, the NOP23 tumor grew uncontrolled (**Fig. S2B**). Blocking lymphocyte egress from secondary lymphoid organs with the sphingosine-1-phosphate receptor (S1PR1) inhibitor FTY720 (25) also abrogated spontaneous tumor control (**Fig. S2C**). It reduced intra-tumor infiltration of lymphocytes as compared to control animals or to mice depleted of CTLs or NK1.1^+^ cells (**Fig. S2D**). Taken together, these results suggested that two-way traffic of immune cells between the periphery and TdLNs was critical for the establishment of an effective endogenous anti-tumor immune response within the TME.

### cDC1 interact with CD4^+^ T cells and tumor-specific CTLs in the TME

Because cDC1, CTL and CD4^+^ T_conv_ were all critical to trigger NOP23 rejection, we wondered whether cDC1 interacted with T cells in the TME. To follow the behavior of Ag-specific CTLs in our experimental models, 1d before tumor engraftment we adoptively transferred low numbers (1,000) of naïve GFP-expressing OT-I cells into *Xcr1^Cre/wt^;Rosa26^tdRFP/wt^* mice that allow in situ visualization of cDC1 by microscopy due to their specific expression of the RFP fluorescent reporter protein (**Fig. S3**). At d7 in the tumor bed, RFP^+^ cDC1 were engaged in cell-cell contacts with both CD4^+^ T cells and anti-tumor CTLs, in close proximity to the HER2^+^ NOP23 cells (**Fig. 3**). This suggested that, in the tumor stroma, cDC1 simultaneously cross-presented tumor Ags to CTLs and relayed them the CD4^+^ T_conv_ help.

**Figure 3.**
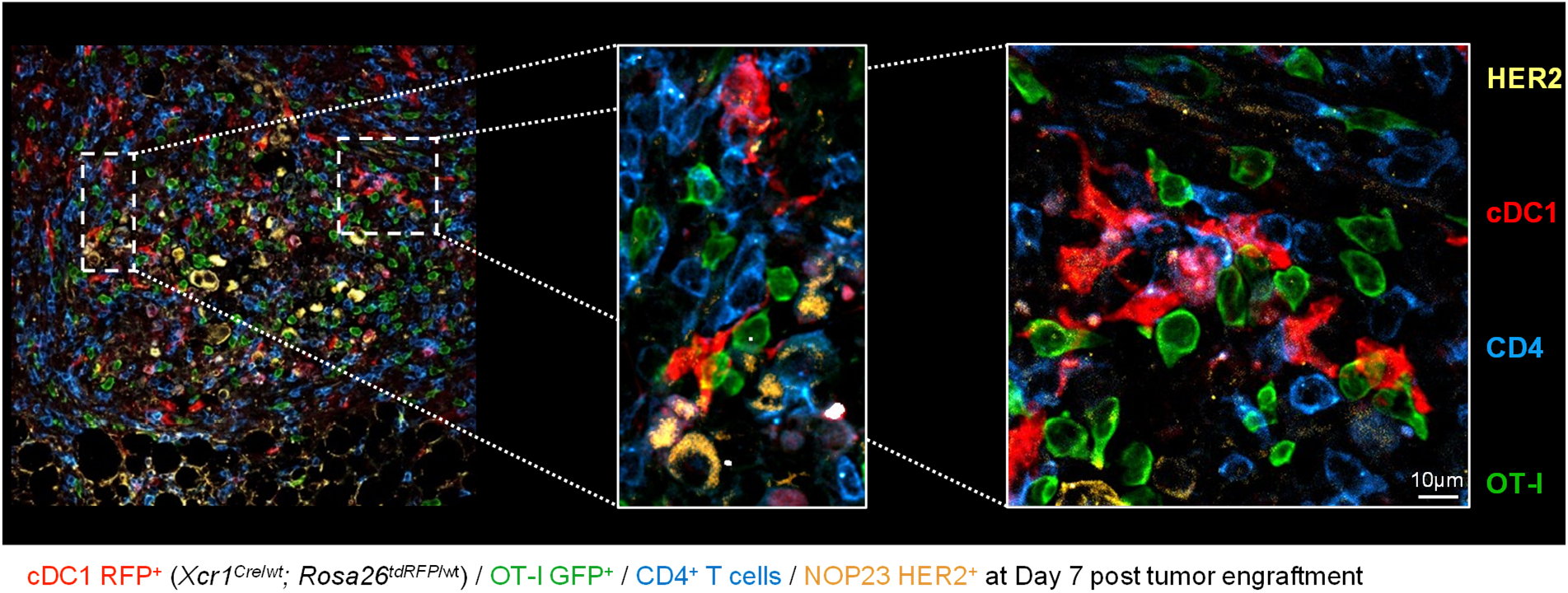
cDC1 interact with CD4^+^ T and tumor-specific CD8^+^ T cells together in the tumor microenvironment. *Xcr1^Cre/wt^;Rosa26^tdRFP/wt^* mice were adoptively transferred with 1,000 GFP^+^ OT-I cells 1d prior to tumor engraftment. 7d post-engraftment, tumors sections were stained for RFP expression (cDC1), GFP (OT-I), CD4 (CD4^+^ T cells) and HER2 (NOP23 cells). This image is representative of 5 individual mice.

### CXCL9 and IL-12 production by cDC1, as well as their trans-presentation of IL-15, are individually dispensable for the immune control of NOP23 tumors

To dissect how cDC1 promoted spontaneous immune control of NOP23, we first tested the candidate molecules CXCL9, IL-12 and IL-15, which had been proposed to be key output signals delivered by cDC1 to T or NK/NK T cells for their recruitment in the tumor or the activation of their effector functions (5). The NOP23 tumors were controlled in *Cxcl9^−/−^, Il12b^−/−^* and *Il15ra* mice as efficiently as in control mice (**Fig. 4A**). Consistent with these results, conditional inactivation of *Cxcl9* or *Il15ra* in cDC1 had no impact on tumor growth (**Fig. S4**). Thus, CXCL9 or IL-12 production and IL-15 transpresentation by cDC1 were not required individually for breast cancer rejection in our experimental settings.

**Figure 4.**
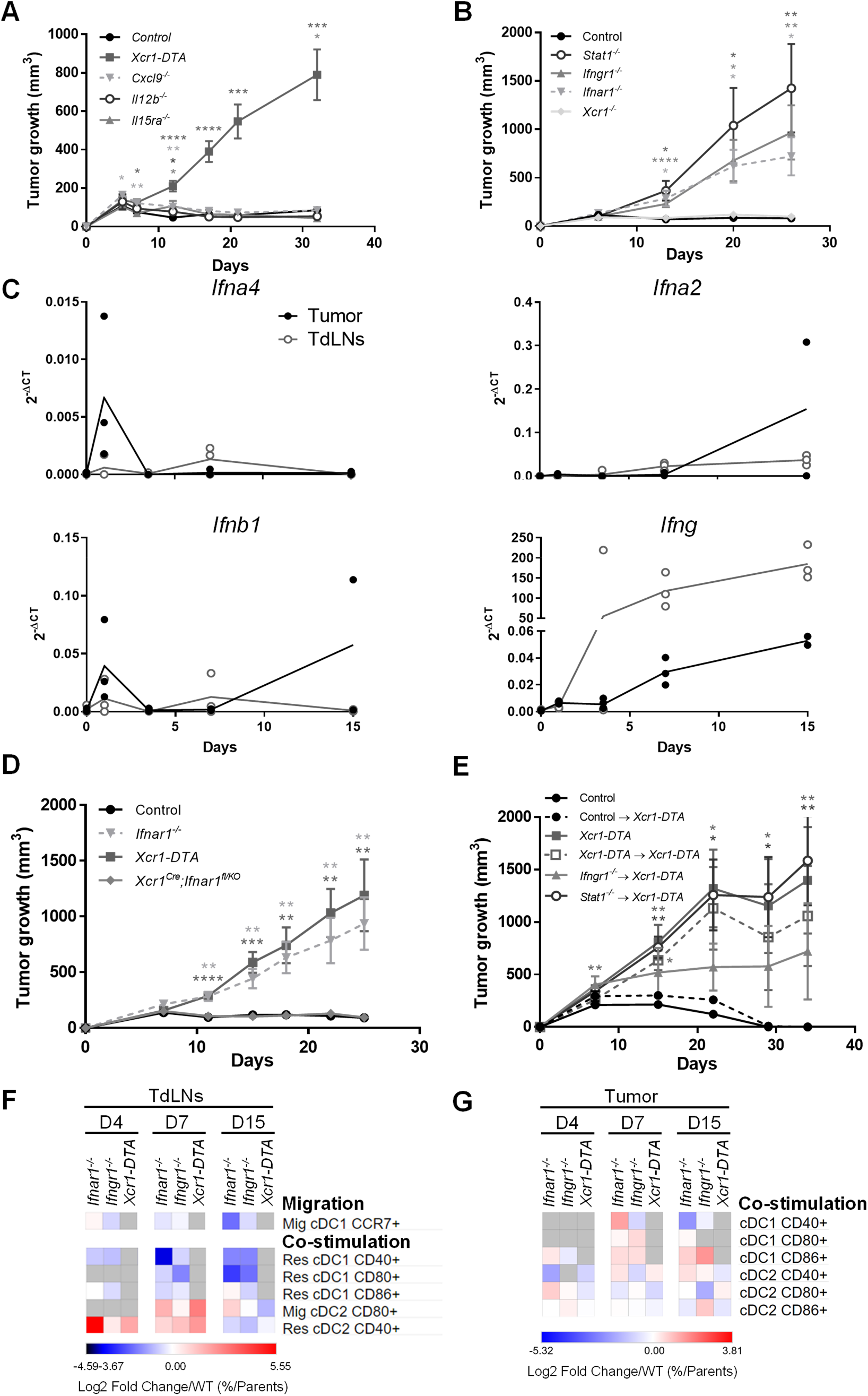
IFNs and cDC1-intrinsic IFN-γ and STAT1 signaling are necessary for breast cancer spontaneous rejection. **A**, Tumor growth in *Xcr1-DTA* (n=10), *Cxcl9^−/−^* (n=9), *Il12b^−/−^* (n=6), *Il15ra^−/−^* (n=5) and control (n=8) female mice. One experiment representative of two independent ones is shown. **B**, Tumor growth in *Xcr1^−/−^* (n=4), *Ifnar1^−/−^* (n=7), *Ifngr1^−/−^* (n=5), *Stat1^−/−^* (n=5) and control (n=6) female mice. One experiment representative of at least 2 independent ones is shown. **C**, Expression analysis of the *Ifna4, Ifna2, Ifnb* and *Ifng* genes in control tumors and their TdLNs (n=2-4) by qPCR. **D**, Tumor growth in *Xcr1-DTA* (n=7), *Ifnar1^−/−^* (n=8), *Xcr1^Cre^;Ifnar1^fl/KO^* (n=13) and control (n=7) female mice. One experiment representative of two independent ones is shown. **E**, Tumor growth in different types of shield bone marrow chimera female mice, control→*Xcr1-DTA* (n=4), *Xcr1-DTA→Xcr1-DTA* (n=4), *Ifngr1^−/−^→Xcr1-DTA* (n=4) and *Stat1^−/−^→Xcr1-DTA* (n=4). *Xcr1-DTA* (n=5) and control (n=5) female mice were used as controls. One experiment representative of two independent ones is shown. *, P < 0.05; **, P < 0.01; ***, P < 0.001; ****, P < 0.0001 (unpaired *t*-test). **F-G**, Heatmaps representing the expression of co-stimulatory receptors on lymphoid resident (res) and migratory (Mig) cDC1 and cDC2 in the TdLNs (**F**) and tumors (**G**) at d4, d7 and d15 after engraftment in *Ifnar1^−/−^, Ifngr1^−/−^, Xcr1-DTA* compared to control mice. The data are shown as Log2 Fold Changes in the ratio of % Population/Parent population from mutant animals to WT (n=3-6 mice per group). The data shown are from two independent experiments pooled together.

### IFN-I and IFN-γ responses, but not XCR1, are necessary for NOP23 tumor control

We next sought to identify the input signals received by cDC1 and promoting their anti-tumor functions. *Xcr1^−/−^* mice efficiently controlled the NOP23 tumor cells (**Fig. 4A**). Hence, XCR1-dependent recruitment of cDC1 to the tumor bed (16) or micro-anatomical attraction to XCL1-producing effector lymphocytes within the tissue (23) was not necessary for breast tumor elimination in our experimental settings. To assess functionally the importance of IFN-I and IFN-γ signaling in NOP23 control, we compared tumor growth between WT animals and *Ifnar1^−/−^, Ifngr1^−/−^* or *Stat1^−/−^* mice, respectively lacking the ability to respond to IFN-I, to IFN-γ or to all types of IFNs including type III IFNs (IFN-III). All three mutant mice failed to control tumor growth, with a more pronounced effect in *Stat1^−/−^* mice (**Fig. 4B**). Thus, both IFN-I and IFN-γ were promoting antitumor immunity in the NOP23 breast cancer model. The analysis of the kinetics of induction of the *Ifna4, Ifna2* and *Ifnb1* genes and of interferon-stimulated genes (ISGs) showed that IFN-I responses were induced in the tumor within the first day after engraftment before rapidly decreasing, while they remained generally low in the TdLNs (**Fig. 4C** and **Fig. S5**). Conversely, *Ifng* expression increased gradually over time and was higher in the TdLNs than in the tumors (**Fig. 4C**). These results suggested that distinct IFNs and ISGs could have complementary roles at different times and locations to initiate and maintain protective anti-tumor immune responses.

### The anti-tumor protective effects of IFN-γ and STAT1 occur at least in part in cDC1, whereas cDC1-intrinsic signaling by IFN-I is dispensable for NOP23 control

We next investigated whether the protective antitumor roles of IFNs were at least in part due to cell-intrinsic effects on cDC1s. NOP23 cells were efficiently rejected in *Xcr1^Cre^;Ifnar1^fl/KO^* and *Karma^Cre/wt^; Ifnar1^fl/fl^* mice that are deficient for IFN-I responsiveness selectively in cDC1 (**Fig. 4D** and **S6A**). Moreover, cDC1 maturation and tumor-specific CTL activation in tumors of *Xcr1^Cre^;Ifnar1^fl/KO^* mice were similar to those in control tumors (**Fig. S6B-C**). Thus, in our experimental settings, cDC1-intrinsic signaling by IFN-I was dispensable for NOP23 control. To determine whether IFN-γ or overall IFN responses in cDC1 were critical for their promotion of NOP23 rejection, we generated shield bone marrow chimera (SBMC) mice deficient selectively in cDC1 for key components of the corresponding signaling pathways, namely *Ifngr1^−/−^→Xcr1-DTA* and *Stat1^−/−^→Xcr1-DTA* animals. Tumors grew progressively in *Ifngr1^−/−^ →Xcr1-DTA* SBMC mice, but to a lower extent than in *Stat1^−/−^→Xcr1-DTA* SBMC mice that harbored a progressive and unabated tumor growth, like *Xcr1-DTA* animals and *Xcr1-DTA→Xcr1-DTA* SBMC mice (**Fig. 4E**). Thus, the beneficial antitumor effects of IFN-γ and STAT1 signaling occurred at least in part in cDC1.

Compared to controls, Mig-cDC1 from *Ifnar1^−/−^* and *Ifngr1^−/−^* TdLNs expressed less CCR7, and their Res-cDC1 were less mature with a decrease in CD40, CD80 and CD86 expression (**Fig. 4F**), contrasting with a higher expression of CD40 on Res-cDC2 at d4-7, and of CD80 on Mig-cDC2 at d7 (**Fig. 4F**). cDC1 expressed less CD40 in the tumors from *Ifnar1^−/−^* and *Ifngr1^−/−^* mice at d15 (**Fig. 4G**). This suggested that loss of IFN responses led to a defective cDC1 maturation in the tumor and TdLNs, with a compensatory increase in cDC2 maturation that was not sufficient to maintain protective antitumor immunity.

### cDC1 and IFNs promote the infiltration of the tumor and its draining lymph node by protective over putatively deleterious immune cell types

To better understand the respective roles of cDC1 and IFNs in the anti-tumor response, we quantified different immune populations in control, *Ifnar1^−/−^, Ifngr1^−/−^* and *Xcr1-DTA* tumor at d4, d7 and d15 (**Fig. 5** and **S7A**). The overall immune cell infiltration in the tumors increased between d4 and d7, the highest in WT mice as compared to mutant animals, and later decreased in all mice (**Fig. 5A**). At d4, the major difference observed between mutant mice and WT controls was a decrease in cDC1 (**Fig. 5B**). At this stage, the tumor was mainly infiltrated in all mouse strains by neutrophils, macrophages and γδ T lymphocytes identified as CD8^-^ CD4^-^ CD3ε^+^ cells (**Fig. S7A**). At day 7, the proportion of cDC1 and NK cells within immune cells remained lower in mutant animals as compared to WT mice (**Fig. 5B-C** and **S7A**). In contrast, the immune infiltrate from the tumors of mutant animals harbored increased proportion of Lin^-^ Siglec-H^-^ CD64^-^ CD11c^+^ MHC-II^-^ cells (**Fig. 5D** and **S7A**), corresponding most likely to DC precursors (26) or immature DC. As compared to WT controls, the mutant mice showed a slight decrease in CD8^+^ T cell proportion at d7 (**Fig. 5D-E** and **S7**), and a marked decrease in the proportions of CD4^+^ T cells mostly at d7 (**Fig. 5D, F** and **S7**). The delayed tumor growth observed upon NK1.1^+^ cell depletion as compared to CTL depletion (**Fig. 1A**) was consistent with the late infiltration of the tumor by NK cells (**Fig. 5C**) and NK T cells (**Fig. 5G**) as compared to T cells (**Fig. 5E-F).** The fraction of activated (CD44^+^) CTLs in the immune infiltrates remained lower in all mutant animals (**Fig. 5D** and **S7**). This was also the case for the fraction of CD4^+^ T cells in *Xcr1-DTA* mice (**Fig. 5D, F** and **S7**), and for the fraction of NK cells in *Ifngr1^−/−^* animals (**Fig. 5C, D** and **S7**). Conversely, at d15, the proportion of neutrophils in the immune infiltrates was much higher in mutant mice than in WT controls (**Fig. 5D**), even though the absolute numbers of neutrophils in tumors decreased sharply over time (**Fig. S7A**). In the TdLNs, the proportion of activated (CD44^+^) CTL was lower at all times in *Xcr1-DTA* and *Ifnar1^−/−^* mice (**Fig. S7B**). Neutrophils were increased over time in all mutant mice, and monocytes or macrophages in *Xcr1*-DTA and *Ifngr1^−/−^* mice, as compared to WT animals (**Fig. S7B**). Starting at d4, Mig-cDC1 accumulated less in *Ifnar1^−/−^* and *Ifngr1^−/−^* mice than in WT animals (**Fig. S7B**). Conversely, the proportion of Mig-cDC2 increased over time (**Fig. S7B**). This suggested that the early migration of cDC1 from the tumor to the TdLNs is IFN-dependent and that its absence leads to a compensatory phenomenon of increased cDC2 migration from the tumor to the TdLN.

**Figure 5.**
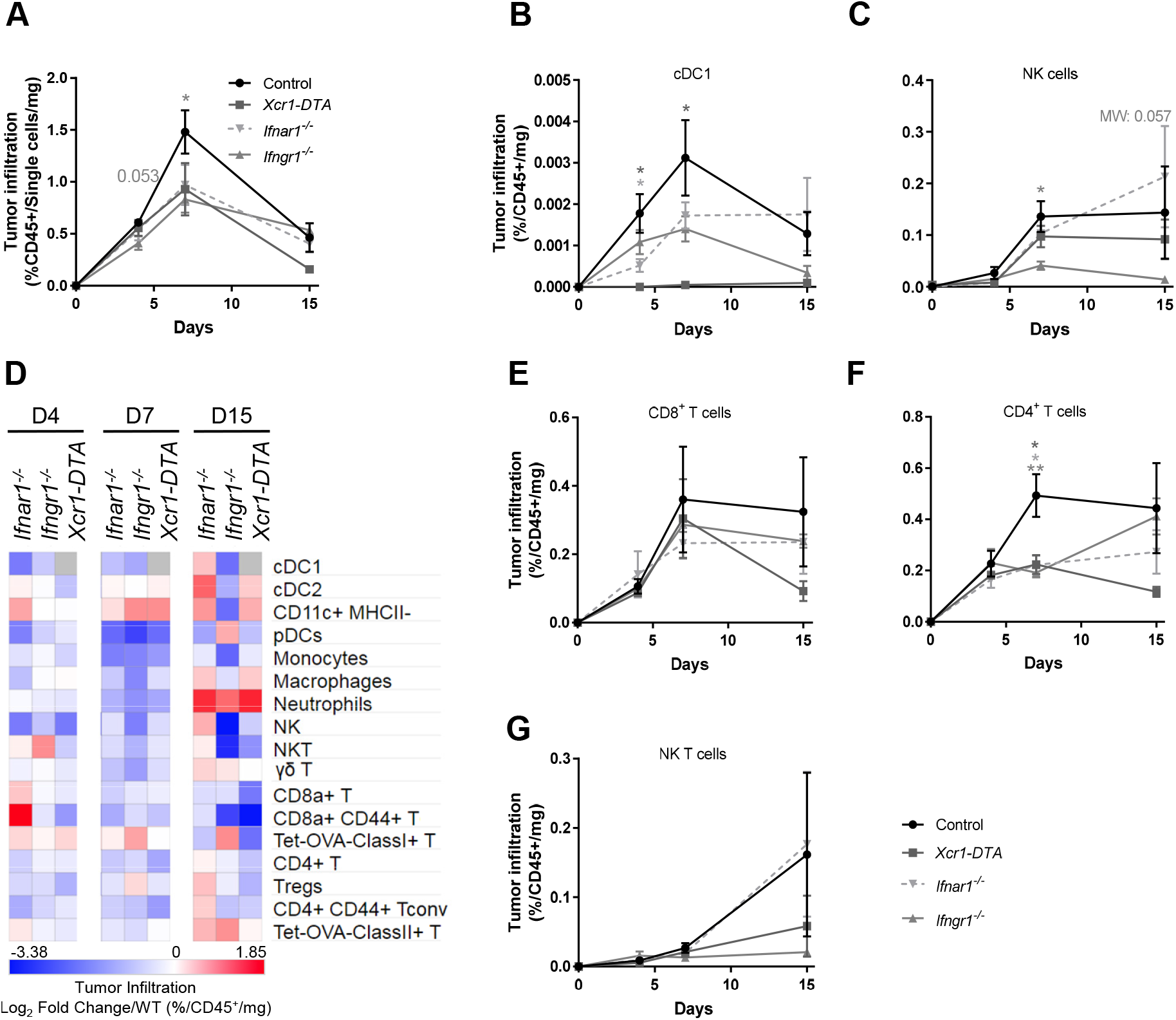
cDC1, type I IFN and type II IFN signaling shape the tumor immune landscape. **A**, Kinetics of the tumor infiltration by CD45^+^ cells at d4, d7 and d15, in control, *Ifnar1^−/−^, Ifngr1^−/−^* and *Xcr1-DTA* mice, as assessed by flow cytometry. **B-C**, Kinetics of the tumor infiltration by cDC1 (B) and NK cells (D) at d4, d7 and d15, in control, *Ifnar1^−/−^, Ifngr1^−/−^* and *Xcr1-DTA* mice. **D**, Heatmap representing the tumor immune landscapes in *Ifnar1^−/−^, Ifngr1^−/−^* and *Xcr1-DTA* mice at d4, d7 and d15. The data are shown as Log2 Fold Changes calculated as the ratio of % Population/CD45^+^/mg of tumor from mutant animals to WT (n=3-6 mice per group). The data shown are from two independent experiments pooled together. **E-G,** Kinetics of the tumor infiltration by CD8^+^ T cells (**E**), CD4^+^ T cells (**F**) and NK T cells (**G**) in control, *Ifnar1^−/−^, Ifngr1^−/−^* and *Xcr1-DTA* mice. For (A-C) and (E-G), the data shown (mean+/−SEM) are from two independent experiments pooled together (n=3-6 mice per group). ns, not significant (p > 0.05); *, p < 0.05; **, p < 0.01; ***, p < 0.001 according to unpaired *t* test or nonparametric Mann-Whitney test (MW) when specified.

Altogether, these results showed that IFNs and cDC1 contributed to sculpt the immune composition of the tumor and TdLN by promoting a higher ratio of protective immune cells, not only NK and CTL but also CD4^+^ T_conv_, over potentially deleterious myeloid cells including macrophages and neutrophils. IFN effects on the tumor immune infiltration may have occurred in part indirectly through promoting early cDC1 recruitment.

### cDC1 and IFNs are necessary for CD4^+^ T_conv_ and CTL terminal activation and effector functions in the TME

We then focused on tumor-specific (Tetramer^+^) CD4^+^ T cells and CTL responses. Their proportions within tumor-infiltrating immune cells did not differ between experimental groups (**Fig. 6A**). As compared to total T cells, tumor-infiltrating Tetramer^+^ T cells expressed higher levels of the checkpoint receptors PD-1, Tim-3 or LAG3 (**Fig. 6B** and **S8**), whose individual expression has been shown to peak at maximal effector phase and reflect CTL activation rather than exhaustion (27,28). Tumor-infiltrating Tetramer^+^ CD4^+^ T cells or CTLs harbored decreased percentages of PD-1^+^ or Tim-3^+^ cells in mutant mice as compared to control animals, suggesting an incomplete effector differentiation (**Fig. 6B** and **Fig. S8**). This was also the case for Tetramer^+^ CTLs in TdLN (**Fig. S9**). The co-expression on the same cell of multiple checkpoint receptors, such as PD-1, Tim-3 and LAG3, has been proposed to define functionally exhausted or dysfunctional lymphocytes (27). The proportion of triple-positive cells were very low on both total and Tetramer^+^ T lymphocytes in all mouse strains, suggesting that the vast majority of anti-tumor T cells were not exhausted (**Fig. S10**). Altogether, these results showed that cDC1 and IFNs were essential to promote CTL and CD4^+^ T cell effector differentiation in the tumor and TdLNs.

We next compared mouse strains for the ability of their tumor- or TdLN-associated T cells to expand and to produce cytokines upon *ex vivo* Ag-specific re-stimulation. Tumor-infiltrating naïve (CD44^-^) and activated (CD44^+^) T_conv_ and CTLs proliferated less in mutant mice than in control animals (**Fig. 6B**). Activated (CD44^+^) CTLs expressed less Granzyme B, IFN-γ and TNF in mutant mice than in control animals, at all-time points examined in the tumor (**Fig. 6B**) and at day 7 in TdLN (**Fig. 6C**). Altogether, these results showed that IFNs and cDC1 in tumor and TdLN contributed to promote the terminal differentiation of anti-tumor CTLs and CD4^+^ T_conv_ for the acquisition of protective effector functions.

### Genes associated to cDC1, CTLs, helper T cells, IFN-I and IFN-γ are associated with a better prognosis in human breast cancer patients

We wanted to know whether the immune cells and signaling pathways associated to the immune control of the mouse NOP23 breast adenocarcinoma model were of good prognosis in breast cancer patients. We used the TCGA database to analyze the transcriptomes of human tumors to infer the degree of their infiltration by cDC1 and its association with the clinical outcome. XCR1 is the only marker fully specific for cDC1 in both mice and humans (17,29). The patients who harbored a high *XCR1* expression in their tumor had a significantly better overall survival (**Fig. 7A**), strongly suggesting that a higher cDC1 infiltration in breast tumors was associated with a better prognosis. Patients whose tumor harbored a higher expression of *CCR7* and of its ligand *CCL19* also had a significantly better overall survival (**Fig. 7B**), suggesting that activation of the CCR7/CCL19 axis in human breast tumors promotes more efficient immune responses as shown in the NOP23 mouse model. Finally, we used gene ontology (GO) to determine whether the gene lists associated with a good (n=367, **Fig. 7C**) or bad (n=210, **Fig. S11**) prognosis in breast cancer patients were enriched for associations with specific biological processes or signaling pathways. The GO terms enriched in the gene list associated to a good prognosis were linked to the activation and helper or cytotoxic functions of T cell responses as well as to IFN-I and IFN-γ signaling pathways (**Fig. 7C**). Conversely, the GO terms associated to the bad prognosis gene list were linked to mitochondria and translation, likely reflecting active metabolism and proliferation of tumor cells (**Fig. S11**). In conclusion, these analyses strongly suggested that cDC1, CD4^+^ T cells, CTLs, NK cells and IFNs play together a crucial role in the immune control of breast cancer, not only in the NOP23 mouse model but also in human patients.

**Figure 6.**
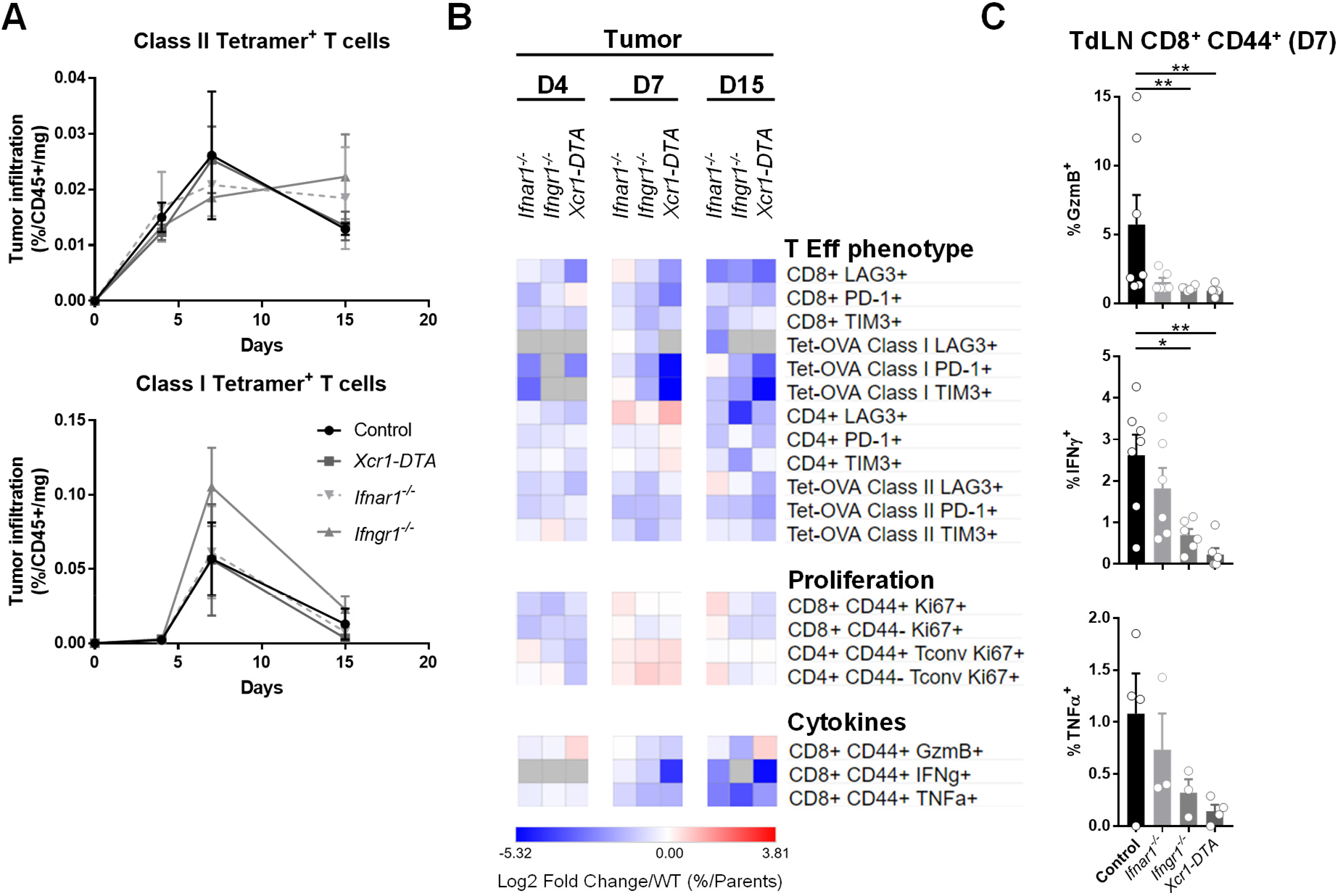
cDC1, type I IFN and type II IFN signaling are necessary for CD4^+^ and CD8^+^ T cell terminal activation and effector functions in the TME. **A**, Kinetics of the tumor infiltration by Ag-specific CD4^+^ (up) and CD8^+^ (bottom) T cells at d4, d7 and d15 in *Ifnar1^−/−^, Ifngr1^−/−^Xcr1-DTA* and control mice. The data shown (mean+/−SEM) are from two independent experiments pooled together (n=3-6 mice per group). **B**, Heatmap representing T cell effector phenotype and proliferation, and CTL cytokine production. The data are shown as Log2 Fold Changes in the ratio of % Population/Parent population from mutant animals to WT (n=3-6 mice per group). **C**, Expression of GzmB, IFN-γ and TNFα by OVA-specific CD44^+^ CD8^+^ T cells in the TdLN of *Ifnar1^−/−^, Ifngr1^−/−^ Xcr1-DTA* and control mice day 7 post-engraftment. The data shown are from two independent experiments pooled together. *, p < 0.05; **, p < 0.01; ***, p < 0.001; (non-parametric Mann-Whitney test).

**Figure 7.**
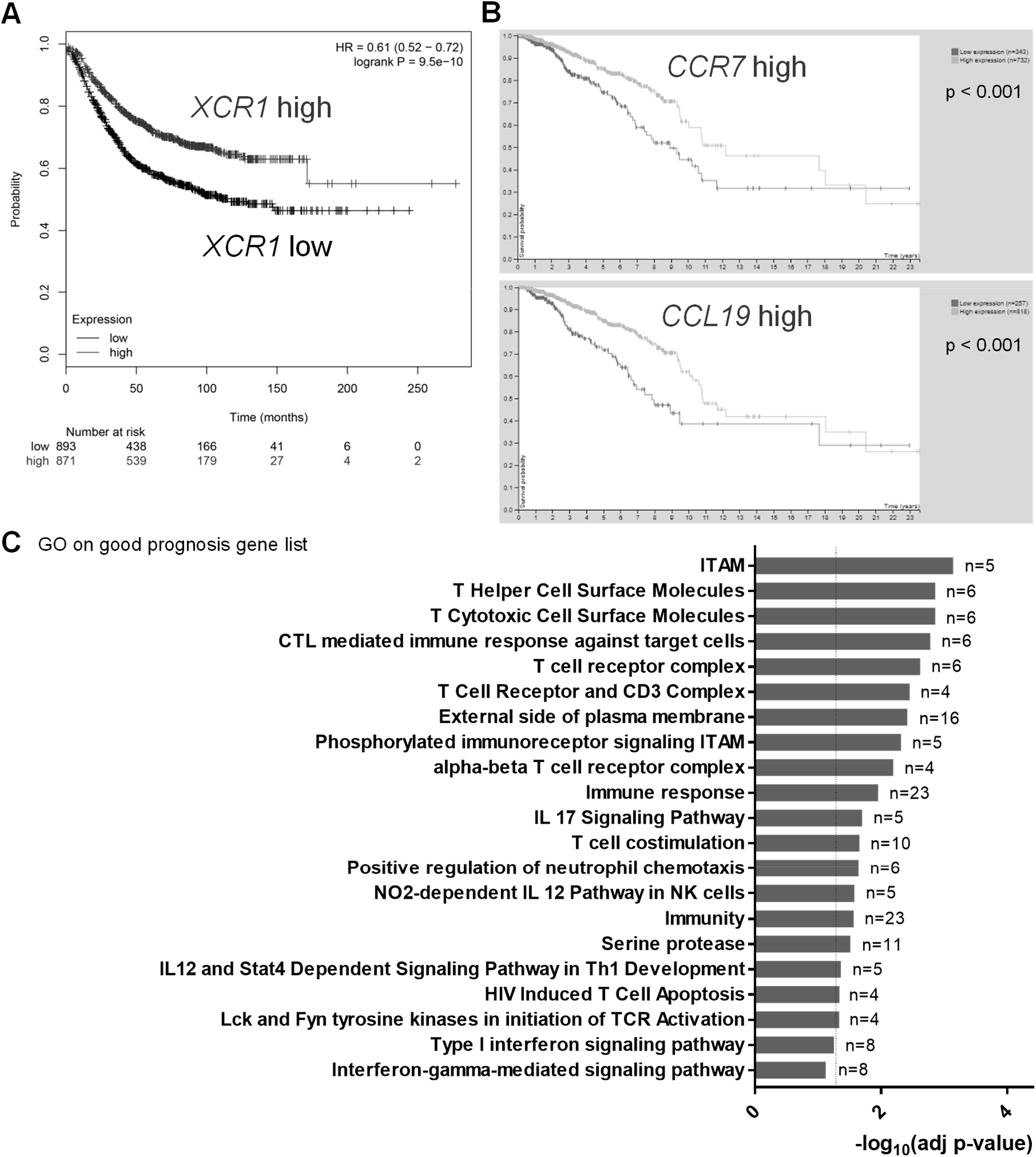
Intratumor expression of *XCR1* and gene ontology annotations linked to CTL, Helper T cell and IFN-I/II signaling are all associated to a better prognosis in human breast cancer patients. **A-B**, Kaplan Meier plot of *XCR1* expression (**A**), and of *CCR7* and *CCL19* expression (**B**) in human breast cancer. **C,** Gene Ontology of breast cancer good prognosis gene set performed with DAVID 6.8. A gene is considered prognostic if correlation analysis of gene expression and clinical outcome resulted in Kaplan-Meier plots with high significance (p<0.001). (A) is from Kaplan Meier-plotter database, (B) and (C) data from the TCGA database.

## Discussion

Here, we investigated whether and how cDC1 promote the spontaneous control of breast cancer in mice, by combining mutant animals enabling specific targeting of cDC1 *in vivo* with an orthotopic model of engraftment of the syngeneic breast adenocarcinoma cell line NOP23. Specifically, we investigated how tumor growth and the nature of the immune infiltrate in the tumor or its draining lymph node were affected by a specific, constitutive or conditional, depletion of cDC1, or by the genetic inactivation of candidate input or output signals specifically in cDC1.

We showed unequivocally that cDC1 were required for the CTL-dependent control of the NOP23 breast adenocarcinoma, throughout the entire antitumor immune response, but especially very early on at the time when anti-tumor T cells are primed. We then wondered which molecular mechanisms promoted cDC1 infiltration into the tumor, and their later migration to the draining lymph node. CXCL9 has been proposed to promote infiltration of pre-cDC1 in the tumor bed (30). XCR1 expression by cDC1 can contribute to their recruitment by XCL1-producing NK/NK T cells and CTLs in infected tissues (23,31). However, neither XCR1 nor CXCL9 were individually required for this function in the NOP23 model. Hence, different chemokine receptors may redundantly promote cDC1 recruitment into the tumor bed, close to effector lymphocytes, as observed for XCR1 and CCR5 in melanoma or colorectal tumors in mice (16). The lack of tumor control in *Ccr7^−/−^* mice could reflect at least in part a strict requirement of this chemokine receptor for cDC1 migration from the tumor to the TdLNs, as previously suggested in a model of melanoma (32).

We then investigated the role of candidate output signals delivered by cDC1 for the promotion of the recruitment of cytotoxic lymphocytes to the tumor and for the activation of their antitumor functions. In microbial infections, cDC1 production of high levels of CXCL9 promote the recruitment of effector and memory CTLs expressing CXCR3 in secondary lymphoid organs (22). This has been proposed to be also the case in tumors (33–35). cDC1 are also a major source of IL-12 and IL-15 promoting activation, survival and cytotoxic functions of NK cells and CTLs (22,34–40). However, CXCL9 and IL-12 production as well as IL-15 transpresentation by cDC1 were individually dispensable for the immune control of the NOP23 breast adenocarcinoma. These results suggest a level of redundancy higher than expected between different types of output signals delivered by cDC1. Indeed, CXCL9 and CXCL10 can both promote the recruitment of CXCR3-expressing effector or memory CTLs in inflamed tissues. In our experimental settings, IL-12, IL-15, IL-18 and IFN-I/III or other cytokines might exert overlapping effects for promoting the proliferation, IFN-γ production and cytotoxic activity of NK and CTLs, as had been reported during certain microbial infections (41,42).

Finally, we sought to identify key input signals required for cDC1 to mediate protective antitumor effects. During microbial infections, the induction of protective NK and T cell responses critically depends on the immunogenic maturation of cDC1, which is driven at least in part by their responses to IFN-I (20) or IFN-γ (22,43,44). Here, we showed that overall IFNI, IFN-γ and STAT1 responses were critical for rejection of the NOP23 breast adenocarcinoma. Only IFN-γ and STAT1 responses were required to occur in cDC1, whereas cell-intrinsic response to IFN-I in cDC1 were dispensable for tumor rejection. Hence, we show here that IFN-I is not always critical for enhancing cDC1 cross-presentation and more generally for licensing them to promote tumor control. This raises the question of the extent to which cDC1-intrisic IFN-I signaling is critical for effective immunity against cancer, besides for immune-mediated rejection of syngeneic and immunogenic fibrosarcoma and melanoma in mice (18,19). However, cDC1-specific inactivation of *Stat1* led to a higher tumor growth than the loss of IFN-γ signaling alone, suggesting some level of redundancy between these two activation pathways for the promotion of cDC1 immunogenic maturation. STAT1 is also key in transducing the signal of IFN-III, whose expression is associated with a better prognosis for breast cancer patients, together with the level of cDC1 infiltration in the tumor (45). Moreover, cDC1 are a main source of IFN-III in human breast tumors (45). Hence, it would be interesting in future studies to investigate whether cDC1 produce, and respond to, IFN-III in the NOP23 breast adenocarcinoma model, and how this may contribute to the spontaneous rejection of the tumor. It will also be interesting to investigate whether further boosting cDC1 development and IFN-III production could help improve immune responses against triple negative breast cancer, similarly to what was recently reported in a therapeutic vaccination trial in human melanoma patients (46). Since overall IFN-I responses but not cDC1 responses to IFN-I were essential for the immune control of the NOP23 breast adenocarcinoma, IFN-I must exert critical effect on other immune cells, likely CTL themselves as was reported in microbial infections (42,47,48).

Finally, to attempt better understanding how IFN and cDC1 were promoting immune rejection of the NOP23 breast adenocarcinoma, we examined how their loss affected the immune landscape of the tumor bed and of the TdLN. We observed that cDC1 and IFNs shaped the tumor immune landscape by promoting CD4^+^ T_conv_ and CTL infiltration and their terminal differentiation with enhanced effector functions. Conversely, cDC1 and IFN responses limited the numbers of putatively deleterious myeloid cell types in the tumor and in TdLN, including, macrophages and neutrophils. In the tumors from *Ifnar1^−/−^* and *Ifngr1^−/−^* mice, cDC1 expressed less CD40. Together with our observation of the simultaneous interactions of cDC1 with CD4^+^ T cells and CTLs in the tumor bed, this suggested that cell-intrinsic responses of cDC1 to IFN may be critical to promote their ability to deliver to CTL the help from CD4^+^ T_conv_ in a manner depending on their interactions via CD40/CD40L. This hypothesis is consistent with the demonstration that simultaneous presentation of viral antigens by cDC1s to CTLs and CD4^+^ T cells is key for robust antiviral cellular immunity (49,50) and with publications linking CD40 expression on cDC1, their ability to activate CD4^+^ T_conv_ and the CTL-dependent rejection of tumors (14,51).

To the best of our knowledge, we show here for the first time in tumors that cDC1 act as a unique cellular platform docking simultaneously CD4^+^ T_conv_ and CTLs, which is likely key to their ability to relay CD4^+^ T cell help to CTLs in situ in tumor, akin to what had been previously shown in infectious settings (49,50). We also rigorously demonstrate for the first time that cell-intrinsic signaling by IFN-γ and STAT1 in cDC1 is critical to their anti-tumor functions. Although this had been suggested before (19,52), it was never formally proven due to lack of adequate models to specifically inactivate IFN-γ signaling in cDC1 without also affecting it in other CD11c^+^ cells. Moreover, in our experimental settings, the licensing of cDC1 by IFN-γ for promoting the control of the NOP23 breast cancer model does not require IL-12 contrary to what has been previously observed or proposed (36,40,52). Beside tumor Ag crosspresentation, the precise nature of the output signals delivered by cDC1 that are critical to induce and maintain protective functions of antitumor CD4^+^ T_conv_ and CTLs remain to be identified, but may encompass IFN-I/III production and CD40 expression. Future studies based on comparative gene expression profiling of WT versus *Ifngr1^−/−^* or *Stat1^−/−^* cDC1 infiltrating the tumor or having migrating to the TdLN should help to identify the output signals whose delivery from cDC1 to effector anti-tumor lymphocytes is critical for the spontaneous rejection of the NOP23 cancer adenocarcinoma model.

Consistent with our experimental results in mice, human breast cancer patients harboring a high expression in their tumor of genes specific to cDC1, CTL, helper T cells or IFN responses have a significantly better clinical outcome. Therefore, we propose the following model of how cDC1 promote tumor immunosurveillance (**Fig. S12**). Following tumor cell immunogenic death, cDC1 capture tumor Ag, undergo immunogenic maturation and migrate to the TdLNs in a CCR7-dependent manner to prime CD4^+^ T_conv_ towards T_H_1 and CD8^+^ T cells towards multipotent protective CTLs. In turn, activated anti-tumor T cells infiltrate the tumor, contributing i) to enhance local recruitment of cDC1, by producing redundant chemokines such as XCL1 and CCL5, and ii) to induce their immunogenic maturation via IFN-γ. This leads to quadripartite interactions in the tumor between cDC1, tumor cells, CD4^+^ T_conv_ and CTLs, ensuring local amplification and maintenance of the effector functions of anti-tumor T cell responses, leading to tumor eradication. IFN-I and IFN-III produced by the cDC1 themselves or other cells might also redundantly contribute to induce cDC1 immunogenic maturation, together with IFN-γ.

## Authors’ Contributions

**R. Mattiuz:** Conceptualization, methodology, investigation, formal analysis, visualization, writing-original draft, writing-review and editing. **C. Brousse:** Resources, investigation. **M. Ambrosini:** Resources, investigation, visualization. **J.-C. Cancel:** Investigation. **Julie Mussard:** investigation. **Amélien Sanlaville:** Resources. **G. Bessou:** Resources, investigation. **C. Caux:** Resources, funding acquisition, methodology, writing– review and editing. **N. Bendriss-Vermare:** Resources, methodology, writing– review and editing. **J. Valladeau-Guilemond:** Resources, methodology, writing– review and editing. **M. Dalod:** Conceptualization, funding acquisition, methodology, supervision, formal analysis, visualization, data curation, validation, writing– review and editing. **K. Crozat:** Conceptualization, funding acquisition, methodology, supervision, investigation, formal analysis, visualization, data curation, validation, writing– review and editing.

## Acknowledgments

This work was supported by grants from the ANR (XCR1-DirectingCells to KC), the Fondation pour la Recherche Médicale (label Equipe FRM 2011, project number DEQ20110421284 to MD), the Fondation ARC (to KC), the Institut National du Cancer (INCa PLBIO 2018-152 to CC and MD), and from the European Research Council under the European Community’s Seventh Framework Program (FP7/2007–2013 grant agreement number 281225 to MD). This work also benefited from institutional funding by CNRS and INSERM. RM was supported by doctoral fellowships from the Biotrail Ph.D. program (Fondation A*MIDEX) and from Fondation ARC. JCC had a doctoral fellowship from LNCC. We thank Dr. Brad Nelson (Director, Deeley Research Centre, BC Cancer Agency, Victoria BC, Canada) for authorization to use the NOP23 cell line. We thank members of the CIML cytometry, Immagimm and histology facilities and from the CIML/CIPHE mouse houses. We acknowledge France Bio lmaging infrastructure supported by the Agence Nationale de la Recherche (ANR-10-INBS-04-01, call “Investissement d’Avenir”). We thank Hugues Lelouard (CIML) for sharing mice, and Rebecca Gentek and Pierre Golstein (CIML) for helpful discussions and protocols. This work benefited from data assembled by the TCGA, The Human Protein Atlas, and Kaplan Meier-plotter databases. Figure S12 has been created with BioRender (https://biorender.com/) under academic license.

## References

1. Vu Manh TP, Bertho N, Hosmalin A, Schwartz-Cornil I, Dalod M. Investigating Evolutionary Conservation of Dendritic Cell Subset Identity and Functions. Front Immunol 2015;6:260 doi:10.3389/fimmu.2015.00260.

2. Sancho D, Joffre OP, Keller AM, Rogers NC, Martinez D, Hernanz-Falcon P, et al. Identification of a dendritic cell receptor that couples sensing of necrosis to immunity. Nature 2009;458(7240):899–903 doi:10.1038/nature07750.

3. Theisen DJ, Davidson JTt, Briseno CG, Gargaro M, Lauron EJ, Wang Q, et al. WDFY4 is required for cross-presentation in response to viral and tumor antigens. Science 2018;362(6415):694–9 doi:10.1126/science.aat5030.

4. Laoui D, Keirsse J, Morias Y, Van Overmeire E, Geeraerts X, Elkrim Y, et al. The tumour microenvironment harbours ontogenically distinct dendritic cell populations with opposing effects on tumour immunity. Nat Commun 2016;7:13720 doi:10.1038/ncomms13720.

5. Cancel JC, Crozat K, Dalod M, Mattiuz R. Are Conventional Type 1 Dendritic Cells Critical for Protective Antitumor Immunity and How? Front Immunol 2019;10:9 doi:10.3389/fimmu.2019.00009.

6. Wculek SK, Cueto FJ, Mujal AM, Melero I, Krummel MF, Sancho D. Dendritic cells in cancer immunology and immunotherapy. Nat Rev Immunol 2020;20(1):7–24 doi:10.1038/s41577-019-0210-z.

7. Sichien D, Scott CL, Martens L, Vanderkerken M, Van Gassen S, Plantinga M, et al. IRF8 Transcription Factor Controls Survival and Function of Terminally Differentiated Conventional and Plasmacytoid Dendritic Cells, Respectively. Immunity 2016;45(3):626–40 doi:10.1016/j.immuni.2016.08.013.

8. Bosteels C, Neyt K, Vanheerswynghels M, van Helden MJ, Sichien D, Debeuf N, et al. Inflammatory Type 2 cDCs Acquire Features of cDC1s and Macrophages to Orchestrate Immunity to Respiratory Virus Infection. Immunity 2020;52(6):1039–56 e9 doi:10.1016/j.immuni.2020.04.005.

9. Hildner K, Edelson BT, Purtha WE, Diamond M, Matsushita H, Kohyama M, et al. Batf3 deficiency reveals a critical role for CD8alpha+ dendritic cells in cytotoxic T cell immunity. Science 2008;322(5904):1097–100 doi:10.1126/science.1164206.

10. Lee W, Kim HS, Hwang SS, Lee GR. The transcription factor Batf3 inhibits the differentiation of regulatory T cells in the periphery. Exp Mol Med 2017;49(11):e393 doi:10.1038/emm.2017.157.

11. Ataide MA, Komander K, Knopper K, Peters AE, Wu H, Eickhoff S, et al. BATF3 programs CD8(+) T cell memory. Nat Immunol 2020;21(11):1397–407 doi:10.1038/s41590-020-0786-2.

12. Qiu Z, Khairallah C, Romanov G, Sheridan BS. Cutting Edge: Batf3 Expression by CD8 T Cells Critically Regulates the Development of Memory Populations. J Immunol 2020;205(4):901–6 doi:10.4049/jimmunol.2000228.

13. Mattiuz R, Wohn C, Ghilas S, Ambrosini M, Alexandre YO, Sanchez C, et al. Novel Cre-Expressing Mouse Strains Permitting to Selectively Track and Edit Type 1 Conventional Dendritic Cells Facilitate Disentangling Their Complexity in vivo. Front Immunol 2018;9:2805 doi:10.3389/fimmu.2018.02805.

14. Ferris ST, Durai V, Wu R, Theisen DJ, Ward JP, Bern MD, et al. cDC1 prime and are licensed by CD4(+) T cells to induce anti-tumour immunity. Nature 2020;584(7822):624–9 doi:10.1038/s41586-020-2611-3.

15. Balan S, Radford KJ, Bhardwaj N. Unexplored horizons of cDC1 in immunity and tolerance. Adv Immunol 2020;148:49–91 doi:10.1016/bs.ai.2020.10.002.

16. Bottcher JP, Bonavita E, Chakravarty P, Blees H, Cabeza-Cabrerizo M, Sammicheli S, et al. NK Cells Stimulate Recruitment of cDC1 into the Tumor Microenvironment Promoting Cancer Immune Control. Cell 2018;172(5):1022–37 e14 doi:10.1016/j.cell.2018.01.004.

17. Crozat K, Guiton R, Contreras V, Feuillet V, Dutertre CA, Ventre E, et al. The XC chemokine receptor 1 is a conserved selective marker of mammalian cells homologous to mouse CD8alpha+ dendritic cells. J Exp Med 2010;207(6):1283–92 doi:10.1084/jem.20100223.

18. Diamond MS, Kinder M, Matsushita H, Mashayekhi M, Dunn GP, Archambault JM, et al. Type I interferon is selectively required by dendritic cells for immune rejection of tumors. J Exp Med 2011;208(10):1989–2003 doi:10.1084/jem.20101158.

19. Fuertes MB, Kacha AK, Kline J, Woo SR, Kranz DM, Murphy KM, et al. Host type I IFN signals are required for antitumor CD8+ T cell responses through CD8{alpha}+ dendritic cells. J Exp Med 2011;208(10):2005–16 doi:10.1084/jem.20101159.

20. Tomasello E, Pollet E, Vu Manh TP, Uze G, Dalod M. Harnessing Mechanistic Knowledge on Beneficial Versus Deleterious IFN-I Effects to Design Innovative Immunotherapies Targeting Cytokine Activity to Specific Cell Types. Front Immunol 2014;5:526 doi:10.3389/fimmu.2014.00526.

21. Martin ML, Wall EM, Sandwith E, Girardin A, Milne K, Watson PH, et al. Density of tumour stroma is correlated to outcome after adoptive transfer of CD4+ and CD8+ T cells in a murine mammary carcinoma model. Breast Cancer Res Treat 2010;121(3):753–63 doi:10.1007/s10549-009-0559-y.

22. Alexandre YO, Ghilas S, Sanchez C, Le Bon A, Crozat K, Dalod M. XCR1+ dendritic cells promote memory CD8+ T cell recall upon secondary infections with Listeria monocytogenes or certain viruses. J Exp Med 2016;213(1):75–92 doi:10.1084/jem.20142350.

23. Ghilas S, Ambrosini M, Cancel J-C, Massé M, Lelouard H, Dalod M, et al. NK cells orchestrate splenic cDC1 migration to potentiate antiviral protective CD8+ T cell responses. bioRxiv 2020:2020.04.23.057463 doi:10.1101/2020.04.23.057463.

24. Simpson TR, Li F, Montalvo-Ortiz W, Sepulveda MA, Bergerhoff K, Arce F, et al. Fc-dependent depletion of tumor-infiltrating regulatory T cells co-defines the efficacy of anti-CTLA-4 therapy against melanoma. J Exp Med 2013;210(9):1695–710 doi:10.1084/jem.20130579.

25. Matloubian M, Lo CG, Cinamon G, Lesneski MJ, Xu Y, Brinkmann V, et al. Lymphocyte egress from thymus and peripheral lymphoid organs is dependent on S1P receptor 1. Nature 2004;427(6972):355–60 doi:10.1038/nature02284.

26. Schlitzer A, Sivakamasundari V, Chen J, Sumatoh HR, Schreuder J, Lum J, et al. Identification of cDC1- and cDC2-committed DC progenitors reveals early lineage priming at the common DC progenitor stage in the bone marrow. Nat Immunol 2015;16(7):718–28 doi:10.1038/ni.3200.

27. Blank CU, Haining WN, Held W, Hogan PG, Kallies A, Lugli E, et al. Defining ‘T cell exhaustion’. Nat Rev Immunol 2019;19(11):665–74 doi:10.1038/s41577-019-0221-9.

28. Singer M, Wang C, Cong L, Marjanovic ND, Kowalczyk MS, Zhang H, et al. A Distinct Gene Module for Dysfunction Uncoupled from Activation in Tumor-Infiltrating T Cells. Cell 2016;166(6):1500–11 e9 doi:10.1016/j.cell.2016.08.052.

29. Bachem A, Guttler S, Hartung E, Ebstein F, Schaefer M, Tannert A, et al. Superior antigen cross-presentation and XCR1 expression define human CD11c+CD141+ cells as homologues of mouse CD8+ dendritic cells. J Exp Med 2010;207(6):1273–81 doi:10.1084/jem.20100348.

30. Cook SJ, Lee Q, Wong AC, Spann BC, Vincent JN, Wong JJ, et al. Differential chemokine receptor expression and usage by pre-cDC1 and pre-cDC2. Immunol Cell Biol 2018;96(10):1131–9 doi:10.1111/imcb.12186.

31. Brewitz A, Eickhoff S, Dahling S, Quast T, Bedoui S, Kroczek RA, et al. CD8(+) T Cells Orchestrate pDC-XCR1(+) Dendritic Cell Spatial and Functional Cooperativity to Optimize Priming. Immunity 2017;46(2):205–19 doi:10.1016/j.immuni.2017.01.003.

32. Roberts EW, Broz ML, Binnewies M, Headley MB, Nelson AE, Wolf DM, et al. Critical Role for CD103(+)/CD141(+) Dendritic Cells Bearing CCR7 for Tumor Antigen Trafficking and Priming of T Cell Immunity in Melanoma. Cancer Cell 2016;30(2):324–36 doi:10.1016/j.ccell.2016.06.003.

33. Spranger S, Dai D, Horton B, Gajewski TF. Tumor-Residing Batf3 Dendritic Cells Are Required for Effector T Cell Trafficking and Adoptive T Cell Therapy. Cancer Cell 2017;31(5):711–23 e4 doi:10.1016/j.ccell.2017.04.003.

34. de Mingo Pulido A, Gardner A, Hiebler S, Soliman H, Rugo HS, Krummel MF, et al. TIM-3 Regulates CD103(+) Dendritic Cell Function and Response to Chemotherapy in Breast Cancer. Cancer Cell 2018;33(1):60–74 e6 doi:10.1016/j.ccell.2017.11.019.

35. Bergamaschi C, Pandit H, Nagy BA, Stellas D, Jensen SM, Bear J, et al. Heterodimeric IL-15 delays tumor growth and promotes intratumoral CTL and dendritic cell accumulation by a cytokine network involving XCL1, IFN-gamma, CXCL9 and CXCL10. J Immunother Cancer 2020;8(1) doi:10.1136/jitc-2020-000599.

36. Broz ML, Binnewies M, Boldajipour B, Nelson AE, Pollack JL, Erle DJ, et al. Dissecting the tumor myeloid compartment reveals rare activating antigen-presenting cells critical for T cell immunity. Cancer Cell 2014;26(5):638–52 doi:10.1016/j.ccell.2014.09.007.

37. Beavis PA, Henderson MA, Giuffrida L, Davenport AJ, Petley EV, House IG, et al. Dual PD-1 and CTLA-4 Checkpoint Blockade Promotes Antitumor Immune Responses through CD4(+)Foxp3(-) Cell-Mediated Modulation of CD103(+) Dendritic Cells. Cancer Immunol Res 2018;6(9):1069–81 doi:10.1158/2326-6066.CIR-18-0291.

38. Greyer M, Whitney PG, Stock AT, Davey GM, Tebartz C, Bachem A, et al. T Cell Help Amplifies Innate Signals in CD8(+) DCs for Optimal CD8(+) T Cell Priming. Cell Rep 2016;14(3):586–97 doi:10.1016/j.celrep.2015.12.058.

39. Mittal D, Vijayan D, Putz EM, Aguilera AR, Markey KA, Straube J, et al. Interleukin-12 from CD103(+) Batf3-Dependent Dendritic Cells Required for NK-Cell Suppression of Metastasis. Cancer Immunol Res 2017;5(12):1098–108 doi:10.1158/2326-6066.CIR-17-0341.

40. Ruffell B, Chang-Strachan D, Chan V, Rosenbusch A, Ho CM, Pryer N, et al. Macrophage IL-10 blocks CD8+ T cell-dependent responses to chemotherapy by suppressing IL-12 expression in intratumoral dendritic cells. Cancer Cell 2014;26(5):623–37 doi:10.1016/j.ccell.2014.09.006.

41. Cousens LP, Peterson R, Hsu S, Dorner A, Altman JD, Ahmed R, et al. Two roads diverged: interferon alpha/beta- and interleukin 12-mediated pathways in promoting T cell interferon gamma responses during viral infection. J Exp Med 1999;189(8):1315–28 doi:10.1084/jem.189.8.1315.

42. Thompson LJ, Kolumam GA, Thomas S, Murali-Krishna K. Innate inflammatory signals induced by various pathogens differentially dictate the IFN-I dependence of CD8 T cells for clonal expansion and memory formation. J Immunol 2006;177(3):1746–54 doi:10.4049/jimmunol.177.3.1746.

43. Lee SH, Carrero JA, Uppaluri R, White JM, Archambault JM, Lai KS, et al. Identifying the initiating events of anti-Listeria responses using mice with conditional loss of IFN-gamma receptor subunit 1 (IFNGR1). J Immunol 2013;191(8):4223–34 doi:10.4049/jimmunol.1300910.

44. Deauvieau F, Ollion V, Doffin AC, Achard C, Fonteneau JF, Verronese E, et al. Human natural killer cells promote cross-presentation of tumor cell-derived antigens by dendritic cells. Int J Cancer 2015;136(5):1085–94 doi:10.1002/ijc.29087.

45. Hubert M, Gobbini E, Couillault C, Manh TV, Doffin AC, Berthet J, et al. IFN-III is selectively produced by cDC1 and predicts good clinical outcome in breast cancer. Sci Immunol 2020;5(46) doi:10.1126/sciimmunol.aav3942.

46. Bhardwaj N, Friedlander PA, Pavlick AC, Ernstoff MS, Gastman BR, Hanks BA, et al. Flt3 ligand augments immune responses to anti-DEC-205-NY-ESO-1 vaccine through expansion of dendritic cell subsets. Nature Cancer 2020;1(12):1204–17 doi:10.1038/s43018-020-00143-y.

47. Kolumam GA, Thomas S, Thompson LJ, Sprent J, Murali-Krishna K. Type I interferons act directly on CD8 T cells to allow clonal expansion and memory formation in response to viral infection. J Exp Med 2005;202(5):637–50 doi:10.1084/jem.20050821.

48. Havenar-Daughton C, Kolumam GA, Murali-Krishna K. Cutting Edge: The direct action of type I IFN on CD4 T cells is critical for sustaining clonal expansion in response to a viral but not a bacterial infection. J Immunol 2006;176(6):3315–9 doi:10.4049/jimmunol.176.6.3315.

49. Eickhoff S, Brewitz A, Gerner MY, Klauschen F, Komander K, Hemmi H, et al. Robust Anti-viral Immunity Requires Multiple Distinct T Cell-Dendritic Cell Interactions. Cell 2015;162(6):1322–37 doi:10.1016/j.cell.2015.08.004.

50. Hor JL, whitney PG, Zaid A, Brooks AG, Heath WR, Mueller SN. Spatiotemporally Distinct Interactions with Dendritic Cell Subsets Facilitates CD4+ and CD8+ T Cell Activation to Localized Viral Infection. Immunity 2015;43(3):554–65 doi:10.1016/j.immuni.2015.07.020.

51. Yamauchi T, Hoki T, Oba T, Attwood K, Cao X, Ito F. CD40 and CD80/86 signaling in cDC1s mediate effective neoantigen vaccination and generation of antigen-specific CX3CR1+ CD8+ T cells in mice. bioRxiv 2020:2020.06.15.151787 doi:10.1101/2020.06.15.151787.

52. Garris CS, Arlauckas SP, Kohler RH, Trefny MP, Garren S, Piot C, et al. Successful Anti-PD-1 Cancer Immunotherapy Requires T Cell-Dendritic Cell Crosstalk Involving the Cytokines IFN-gamma and IL-12. Immunity 2018;49(6):1148–61 e7 doi:10.1016/j.immuni.2018.09.024.

