## Supplementary material for "cDC1 and interferons promote spontaneous CD4^+^ and CD8^+^ T cell protective responses to breast cancer": 3 supplementary tables and 12 supplementary figures

**Supplementary Table S1.** Mouse strains used in this study

| Type | **Mouse strains**[1] | **Official nomenclatures** | **References**[2] | **Source** |
|---|---|---|---|---|
| KI | *Karma-tmt-hDTR* | *Gpr141b*$^{tm1(HBEGF)Ciphe}$ | (1) | Dalod lab/Ciphe |
| | *Tg*$^{TcraTcrb1100Mjb}$; *Rag2*$^{-/-}$ | C57BL/6-Tg(TcraTcrb)1100Mjb/J; B6(Cg)-*Rag2*$^{tm1.1Cgn}$/J | (2,3) | CIML: Dr. A.M. Schmidt-Verhulst Jackson Laboratory (Stock n° 003831; 008449) |
| | *Ubc-GFP*$^{+/+}$ | C57BL/6-Tg(UBC-GFP)30Scha/J | (4) | CIML: Dr. M. Bajénoff Jackson Laboratory (Stock n° 004353) |
| KO | *Ccr7*$^{-/-}$ | B6.129P2(C)-*Ccr7*$^{tm1Rfor}$/J | (5) | CIML: Dr. H. Lelouard Jackson Laboratory (Stock n° 006621) |
| | *Cxcl9*$^{-/-}$ | B6-*Cxcl9*$^{tm2Ciphe}$ | This paper | Dalod lab/Ciphe |
| | *Ifnar1*$^{-/-}$ | B6.129S2-*Ifnar1*$^{tm1Agt}$ | (6,7) | Pr. U. Kalinke |
| | *Ifngr1*$^{-/-}$ | B6.129S7-*Ifngr1*$^{tm1Agt}$/J | (8) | CIML: Dr. H. Lelouard Jackson Laboratory (Stock n° 003288) |
| | *Il12b*$^{-/-}$ | B6.129S1-*Il12b*$^{tm1Jm}$/J | (9) | CIML: Dr. H. Lelouard Jackson Laboratory (Stock n° 002693) |
| | *Il15ra*$^{-/-}$ | B6;129X1-*Il15ra*$^{tm1Ama}$/J | (10) | Jackson Laboratory (Stock n° 003723) |
| | *Stat1*$^{-/-}$ | *B6-Stat1*$^{tm1d(EUCOMM)Ciphe}$ | (11) | EUCOMM/ Dalod lab/Ciphe |
| | *Xcr1*$^{-/-}$ | B6.129P2-*Xcr1*$^{tm1Dgen}$/J | (12,13) | Jackson Laboratory (Stock n° 005791) |
| Cre | *Karma*$^{Cre}$ | B6-*Gpr141b*$^{tm2Ciphe}$ | (14) | Dalod lab/Ciphe |
| | *Xcr1*$^{Cre}$ | B6-*Xcr1*$^{tm1Ciphe}$ | (14) | Dalod lab/Ciphe |
| Floxed | *Cxcl9*$^{fl}$ | B6-*Cxcl9*$^{tm1Ciphe}$ | This paper | Dalod lab/Ciphe |
| | *Ifnar1*$^{fl}$ | *Ifnar1*$^{tm1Uka}$ | (15,16) | Pr. U. Kalinke |
| | *Il15ra*$^{fl}$ | C57BL/6-*Il15ra*$^{tm2.1Ama}$/J | (17) | Jackson Laboratory (Stock n° 022365) |
| | *Rosa26*$^{lox-stop-lox\text{-}DTA}$ | B6.129P2-*Gt(ROSA)26Sor*$^{tm1(DTA)Lky}$/J | (18) | Prs. D. Voehringer & R.M. Locksley Jackson Laboratory (Stock n° 009669) |
| | *Rosa26*$^{lox-stop-lox\text{-}hDTR}$ | *Gt(ROSA)26Sor*$^{tm1(HBEGF)Awai}$/J | (19) | CIML: Dr. M. Sieweke Jackson Laboratory (Stock n° 007900) |
| | *Rosa26*$^{lox-stop-lox\text{-}tdRFP}$ | *Gt(ROSA)26Sor*$^{tm1Hjf}$ | (20) | CIML: Dr. H. Luche |

[1] All mice were maintained in a C57BL/6J background.

[2] References:


1. Alexandre YO, Ghilas S, Sanchez C, Le Bon A, Crozat K, Dalod M. XCR1+ dendritic cells promote memory CD8+ T cell recall upon secondary infections with Listeria monocytogenes or certain viruses. J Exp Med **2016**;213(1):75-92 doi 10.1084/jem.20142350.
2. Hao Z, Rajewsky K. Homeostasis of peripheral B cells in the absence of B cell influx from the bone marrow. J Exp Med **2001**;194(8):1151-64 doi 10.1084/jem.194.8.1151.
3. Hogquist KA, Jameson SC, Heath WR, Howard JL, Bevan MJ, Carbone FR. T cell receptor antagonist peptides induce positive selection. Cell **1994**;76(1):17-27 doi 10.1016/0092-8674(94)90169-4.

4. Schaefer BC, Schaefer ML, Kappler JW, Marrack P, Kedl RM. Observation of antigen-dependent CD8+ T-cell/ dendritic cell interactions in vivo. Cell Immunol **2001**;214(2):110-22 doi 10.1006/cimm.2001.1895.
5. Forster R, Schubel A, Breitfeld D, Kremmer E, Renner-Muller I, Wolf E*, et al.* CCR7 coordinates the primary immune response by establishing functional microenvironments in secondary lymphoid organs. Cell **1999**;99(1):23-33 doi 10.1016/s0092-8674(00)80059-8.
6. Muller U, Steinhoff U, Reis LF, Hemmi S, Pavlovic J, Zinkernagel RM*, et al.* Functional role of type I and type II interferons in antiviral defense. Science **1994**;264(5167):1918-21 doi 10.1126/science.8009221.
7. Baranek T, Manh TP, Alexandre Y, Maqbool MA, Cabeza JZ, Tomasello E*, et al.* Differential responses of immune cells to type I interferon contribute to host resistance to viral infection. Cell Host Microbe **2012**;12(4):571-84 doi 10.1016/j.chom.2012.09.002.
8. Huang S, Hendriks W, Althage A, Hemmi S, Bluethmann H, Kamijo R*, et al.* Immune response in mice that lack the interferon-gamma receptor. Science **1993**;259(5102):1742-5 doi 10.1126/science.8456301.
9. Magram J, Connaughton SE, Warrier RR, Carvajal DM, Wu CY, Ferrante J*, et al.* IL-12-deficient mice are defective in IFN gamma production and type 1 cytokine responses. Immunity **1996**;4(5):471-81 doi 10.1016/s1074-7613(00)80413-6.
10. Lodolce JP, Boone DL, Chai S, Swain RE, Dassopoulos T, Trettin S*, et al.* IL-15 receptor maintains lymphoid homeostasis by supporting lymphocyte homing and proliferation. Immunity **1998**;9(5):669-76 doi 10.1016/s1074-7613(00)80664-0.
11. Tomasello E, Naciri K, Chelbi R, Bessou G, Fries A, Gressier E*, et al.* Molecular dissection of plasmacytoid dendritic cell activation in vivo during a viral infection. EMBO J **2018**;37(19) doi 10.15252/embj.201798836.
12. Crozat K, Guiton R, Contreras V, Feuillet V, Dutertre CA, Ventre E*, et al.* The XC chemokine receptor 1 is a conserved selective marker of mammalian cells homologous to mouse CD8alpha+ dendritic cells. J Exp Med **2010**;207(6):1283-92 doi 10.1084/jem.20100223.
13. Dorner BG, Dorner MB, Zhou X, Opitz C, Mora A, Guttler S*, et al.* Selective expression of the chemokine receptor XCR1 on cross-presenting dendritic cells determines cooperation with CD8+ T cells. Immunity **2009**;31(5):823-33 doi 10.1016/j.immuni.2009.08.027.
14. Mattiuz R, Wohn C, Ghilas S, Ambrosini M, Alexandre YO, Sanchez C*, et al.* Novel Cre-Expressing Mouse Strains Permitting to Selectively Track and Edit Type 1 Conventional Dendritic Cells Facilitate Disentangling Their Complexity in vivo. Front Immunol **2018**;9:2805 doi 10.3389/fimmu.2018.02805.
15. Le Bon A, Durand V, Kamphuis E, Thompson C, Bulfone-Paus S, Rossmann C*, et al.* Direct stimulation of T cells by type I IFN enhances the CD8+ T cell response during cross-priming. J Immunol **2006**;176(8):4682-9 doi 10.4049/jimmunol.176.8.4682.
16. Kamphuis E, Junt T, Waibler Z, Forster R, Kalinke U. Type I interferons directly regulate lymphocyte recirculation and cause transient blood lymphopenia. Blood **2006**;108(10):3253-61 doi 10.1182/blood-2006-06-027599.
17. Mortier E, Advincula R, Kim L, Chmura S, Barrera J, Reizis B*, et al.* Macrophage- and dendritic-cell-derived interleukin-15 receptor alpha supports homeostasis of distinct CD8+ T cell subsets. Immunity **2009**;31(5):811-22 doi 10.1016/j.immuni.2009.09.017.
18. Voehringer D, Liang HE, Locksley RM. Homeostasis and effector function of lymphopenia-induced "memory-like" T cells in constitutively T cell-depleted mice. J Immunol **2008**;180(7):4742-53 doi 10.4049/jimmunol.180.7.4742.
19. Buch T, Heppner FL, Tertilt C, Heinen TJ, Kremer M, Wunderlich FT*, et al.* A Cre-inducible diphtheria toxin receptor mediates cell lineage ablation after toxin administration. Nat Methods **2005**;2(6):419-26 doi 10.1038/nmeth762.
20. Luche H, Weber O, Nageswara Rao T, Blum C, Fehling HJ. Faithful activation of an extra-bright red fluorescent protein in "knock-in" Cre-reporter mice ideally suited for lineage tracing studies. Eur J Immunol **2007**;37(1):43-53 doi 10.1002/eji.200636745.

**Supplementary Table S2.** Antibodies used in this study

| Antibody | Clone | Conjugates | Company | Dilution/Dose | Use |
|---|---|---|---|---|---|
| **Arm Hamster** | Polyclonal | A594 | Jackson ImmunoResearch | 1/200 | Microscopy |
| **CD3ε** | 145-2C11 | BV510 | BD Biosciences | 1/200 | Flow cytometry |
| **CD3ε** | 145-2C11 | Purified | BD Biosciences | 1/300 | Flow cytometry |
| **CD4** | GK1.5 | Purified | BioXCell | 500 µg | *In vivo* depletion |
| **CD4** | GK1.5 | APC-H7 | BD Biosciences | 1/200 | Flow cytometry |
| **CD4** | RM4-5 | ef450 | ThermoFisher Scientific | 1/100 | Microscopy |
| **CD8α** | 53-6.7 | PerCP-Cy5.5 | BD Biosciences | 1/200 | Flow cytometry |
| **CD8β** | H35-17.2 | Purified | produced_in_house | 150 µg | *In vivo* depletion |
| **CD11b** | M1/70 | BUV395 | BD Biosciences | 1/400 | Flow cytometry |
| **CD11c** | N418 | BV785 | BioLegend | 1/200 | Flow cytometry |
| **CD19** | 1D3 | BV510 | BD Biosciences | 1/200 | Flow cytometry |
| **CD19** | 1D3 | Alexa700 | BD Biosciences | 1/200 | Flow cytometry |
| **CD24** | M1/69 | eFluor450 | eBioscience | 1/1000 | Flow cytometry |
| **CD25** | PC61 | BV421 | BD Biosciences | 1/600 | Flow cytometry |
| **CD40** | 3/23 | PE | BD Biosciences | 1/200 | Flow cytometry |
| **CD43** | 1B11 | PE-Cy5 | BioLegend | 1/200 | Flow cytometry |
| **CD44** | IM7 | PE-Cy7 | eBioscience | 1/800 | Flow cytometry |
| **CD45.2** | 104 | V500 | BD Biosciences | 1/200 | Flow cytometry |
| **CD45.2** | 104 | BUV737 | BD Biosciences | 1/400 | Flow cytometry |
| **CD62l** | MEL-14 | BV421 | BD Biosciences | 1/400 | Flow cytometry |
| **CD64** | X54-5/7.1 | BV711 | BioLegend | 1/200 | Flow cytometry |
| **CD69** | H1.2F3 | FITC | eBioscience | 1/200 | Flow cytometry |
| **CD80** | 16-10A1 | APC | BD Biosciences | 1/200 | Flow cytometry |
| **CD86** | GL1 | PE-Cy7 | BD Biosciences | 1/400 | Flow cytometry |
| **CD117** | 2B8 | PE-Cy7 | BD Biosciences | 1/200 | Flow cytometry |
| **CD172a** | P84 | FITC | BD Biosciences | 1/200 | Flow cytometry |
| **CCR7** | 4B12 | Biotin | eBioscience | 1/100 | Flow cytometry |
| **CCR7** | 4B12 | PE | BD Biosciences | 1/100 | Flow cytometry |
| **CTLA-4** | 9D9 | Purified | BioXCell | 200 µg | *In vivo* depletion |
| **F4/80** | BM8 | BV605 | BioLegend | 1/200 | Flow cytometry |
| **FcεRIα** | MAR-1 | PacificBlue | BioLegend | 1/200 | Flow cytometry |
| **FoxP3** | FJK-16s | Biotin | eBioscience | 1/100 | Flow cytometry |
| **GFP** | Polyclonal | A488 | Invitrogen | 1/1000 | Microscopy |
| **GzmB** | GB11 | APC | Invitrogen | 1/100 | Flow cytometry |
| **HER2** | 7.16.4 | Purified | BioXCell | 1/1000 | Microscopy |
| **IFNγ** | XMG1.2 | Alexa700 | BD Biosciences | 1/200 | Flow cytometry |
| **Ki67** | B56 | V450 | BD Biosciences | 1/100 | Flow cytometry |
| **LAG3** | C9B7W | BV711 | BD Biosciences | 1/200 | Flow cytometry |
| **Ly-6C** | AL-21 | APC-Cy7 | BD Biosciences | 1/1000 | Flow cytometry |
| **MHC-II** | M5/114.15.2 | Alexa700 | BioLegend | 1/400 | Flow cytometry |
| **NK1.1** | PK136 | Purified | produced_in_house | 200 µg | *In vivo* depletion |
| **NK1.1** | PK136 | BV650 | BioLegend | 1/400 | Flow cytometry |
| **NKp46** | 29A1.4 | BV510 | BD Biosciences | 1/200 | Flow cytometry |
| **PD-1** | 29F.1A12 | BV785 | BioLegend | 1/400 | Flow cytometry |
| **Rabbit** | Polyclonal | A647 | Molecular Probes | 1/500 | Microscopy |
| **RFP** | Polyclonal | Purified | Rockland | 1/500 | Microscopy |
| **Siglec-H** | 551 | PerCP-Cy5.5 | BioLegend | 1/200 | Flow cytometry |
| **TCRβ** | H57-597 | FITC | BD Biosciences | 1/200 | Flow cytometry |
| **Tim-3** | RMT3-23 | BV605 | BioLegend | 1/200 | Flow cytometry |
| **TNFα** | MP6-XT22 | BV785 | BioLegend | 1/200 | Flow cytometry |
| **XCR1** | ZET | BV650 | BioLegend | 1/1000 | Flow cytometry |

**Supplementary Table S3.** Primers used in this study for qRT-PCR

| | Forward primers | Reverse primers |
|---|---|---|
| *Hprt* | 5′-GGCCCTCTGTGTGCTCAAG-3′ | 5′-CTGATAAAATCTACAGTCATAGGAATGGA-3′ |
| *Ifna2* | 5'-AGGACAGGCAGGACTTTGGA-3' | 5'-GCCTTCTGGATCTGCTGGTTA-3' |
| *Ifna4* | 5'-AAGGACAGGAAGGATTTTGGATT-3' | 5'-GAGCCTTCTGGATCTGTTGGTT-3' |
| *Ifnb* | 5'-GGTGGTCCGAGCAGATCTT-3' | 5'-CAGTTTTGGAAGTTTCTGGTAAGTCTT-3' |
| *Ifng* | 5'-CAACAGCAAGGCGAAAAAGG-3' | 5'-CCTGTGGGTTGTTGACCTCAA-3' |
| *Irf7* | 5′-TCCAGTTGATCCGCATAAGGT-3′ | 5′-CTTCCCTATTTTCCGTGGCTG-3′ |
| *Isg15* | 5′-GGTGTCCGTGACTAACTCCAT-3′ | 5′-TGGAAAGGGTAAGACCGTCCT-3′ |
| *Mx1* | 5'-GACCATAGGGGTCTTGACCAA-3' | 5'-AGACTTGCTCTTTCTGAAAAGCC-3' |
| *Oas3* | 5'-TCTGGGGTCGCTAAACATCAC-3' | 5'-GATGACGAGTTCGACATCGGT-3' |

**Supplementary Figure S1**

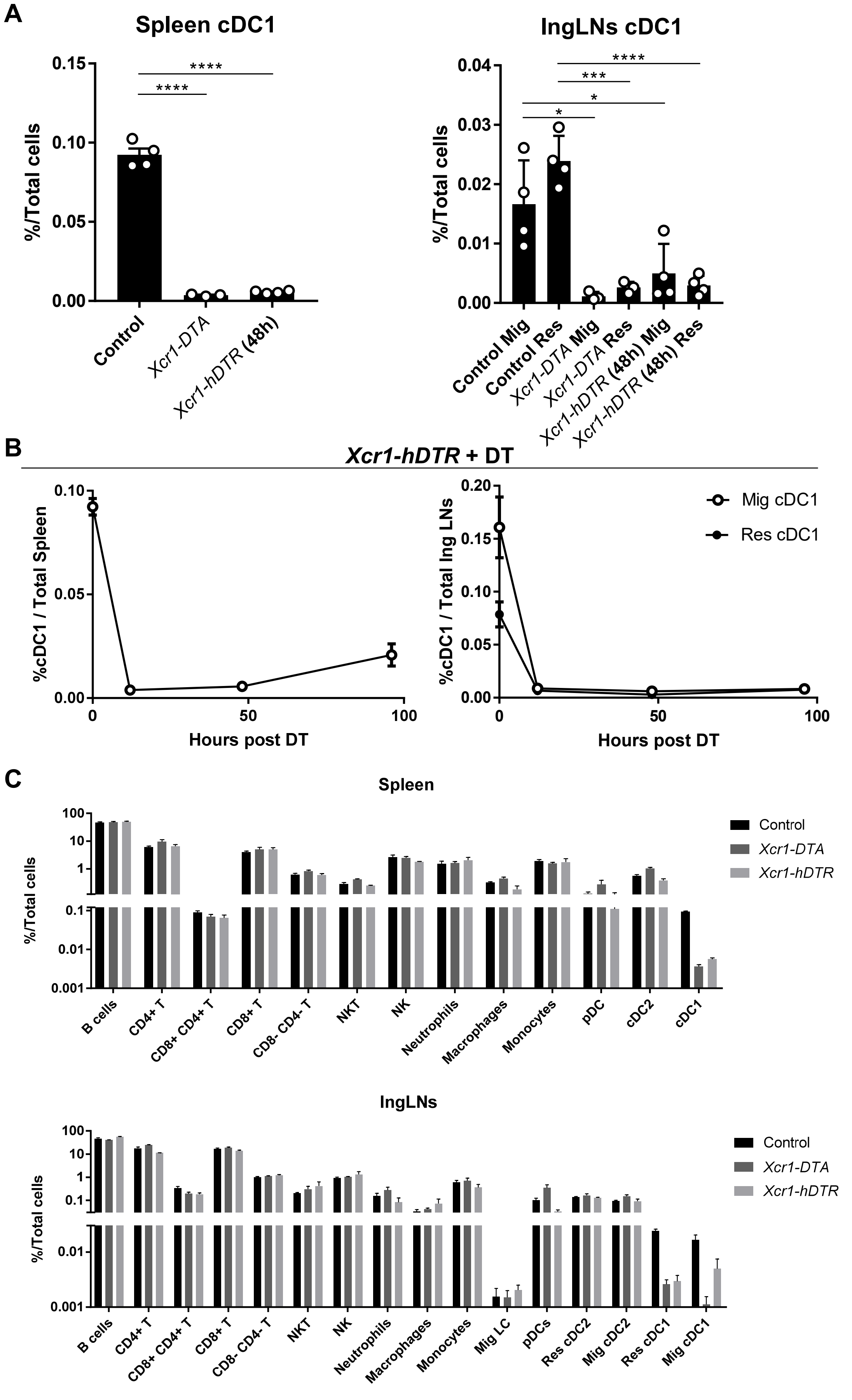


**Supplementary Figure S1.** Efficient cDC1 depletion in *Xcr1-DTA* and *Xcr1-hDTR* mice.

**A,** cDC1 proportion in spleen (left) and IngLNs (right) in *Xcr1-DTA* and *Xcr1-hDTR* mice (48h after DT). **B,** Kinetics of cDC1 depletion in spleen (left) and IngLNs (right) after one DT injection in *Xcr1-*

*hDTR* mice. **C,** %/total cells for the indicated populations in spleen (top) and IngLNs (bottom) in *Xcr1-DTA*, *Xcr1-hDTR* (48h after DT) and control mice. The data shown are for 3-4 mice per condition from one experiment representative of two independent ones. *, P < 0.05; **, P < 0.01; ***, P < 0.001; ****, P < 0.0001 (unpaired *t* test).

### Supplementary Figure S2

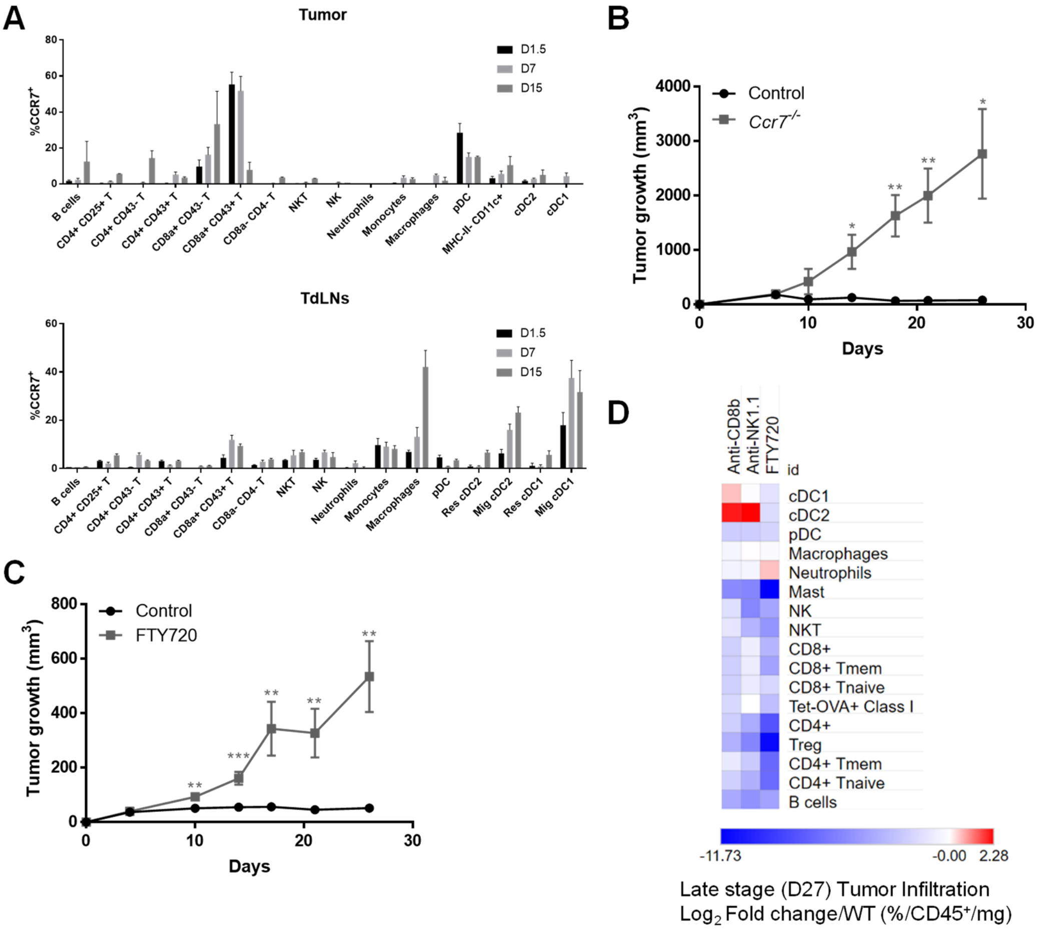


**Supplementary Figure S2.** Trafficking of immune cells into and out of the TdLNs is instrumental in breast cancer spontaneous rejection.

**A,** Kinetics of CCR7 expression among immune populations in control tumor and TdLNs (2-6 mice per time point). **B,** Tumor growth of *Ccr7*$^{-/-}$ (n=3) and control (n=5) females. One experiment representative of two independent ones is shown. **C,** Tumor growth of FTY720-treated (n=7) and untreated (Control, n=7) female mice. One experiment representative of two independent ones is shown. **D,** Heatmap representing the immune landscapes in d27 tumors in CD8$^{+}$ T cell-depleted (anti-CD8β), NK cell-depleted (anti-NK1.1) or in FTY720-treated females compared to Control. The data are shown as Log2 Fold Changes of the %Population/CD45$^{+}$/mg of tumor in analyzed animals compared to WT. n=3-4 mice per group. *, p < 0.05; **, p < 0.01; ***, p < 0.001 (unpaired *t* test).

**Supplementary Figure S3**

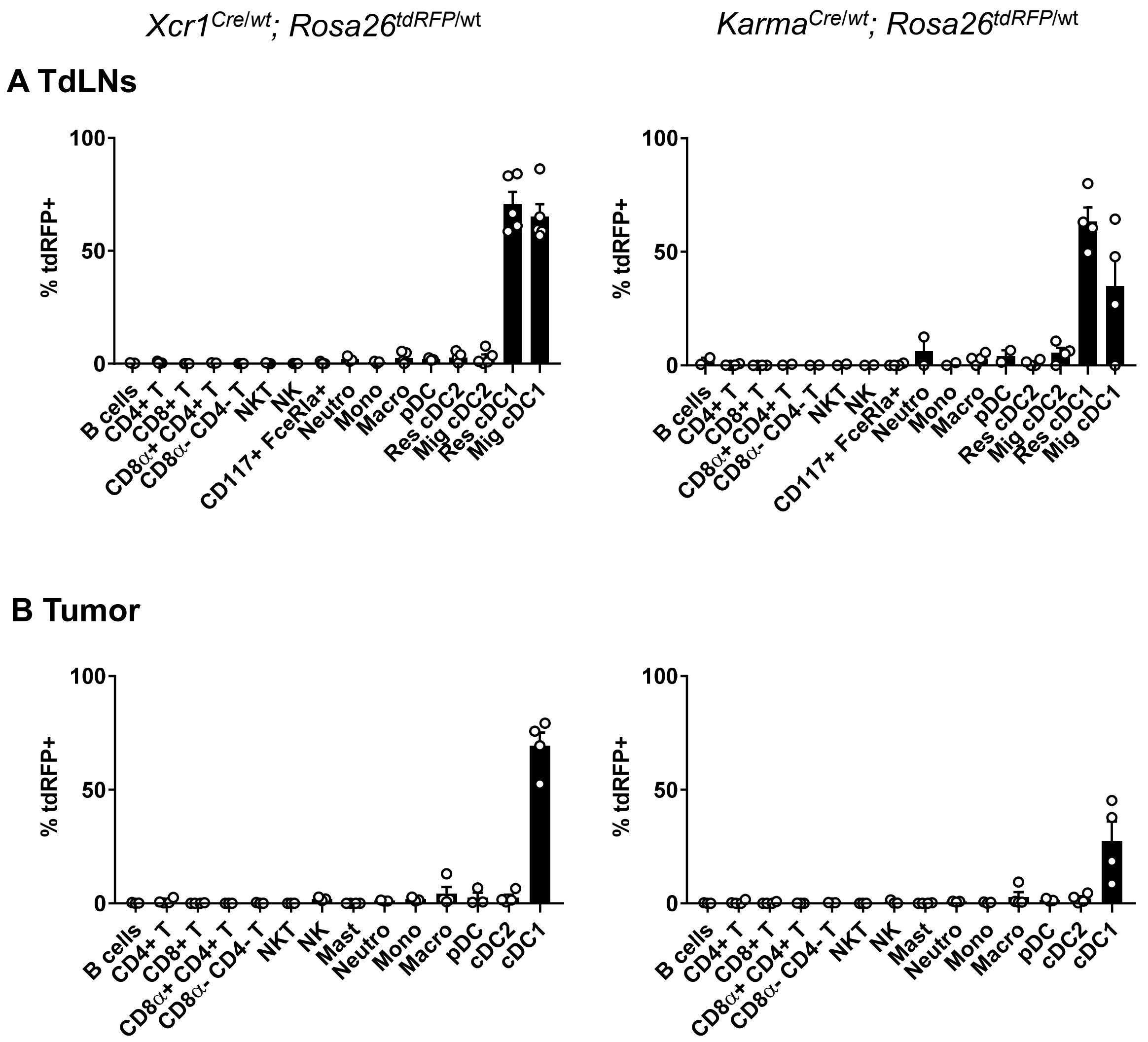


**Supplementary Figure S3.** The *Xcr1*$^{Cre/wt}$*;Rosa26*$^{tdRFP/wt}$ and *Karma*$^{Cre/wt}$*;Rosa26*$^{tdRFP/wt}$ mouse strains allow specific fate mapping of cDC1 in the tumor and TdLNs

tdRFP expression analysis by different immune cell populations from TdLNs (**A**) and tumors (**B**) at d4 post engraftment in *Xcr1*$^{Cre/wt}$*;Rosa26*$^{tdRFP/wt}$ and *Karma*$^{Cre/wt}$*;Rosa26*$^{tdRFP/wt}$ mice. The data shown (one dot per mouse with mean +/- SEM per group) are from two independent experiments pooled together.

### Supplementary Figure S4

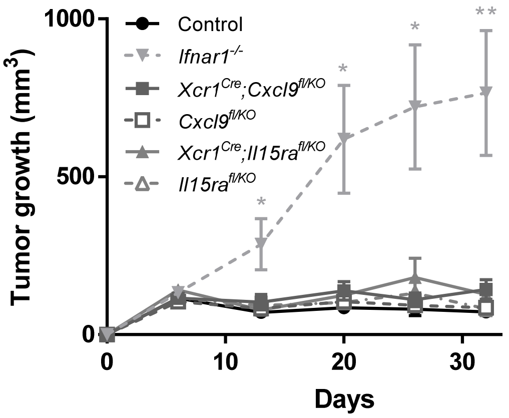


**Supplementary Figure S4.** CXCL9 production and IL-15 trans-presentation by cDC1 are not necessary for tumor rejection.

Tumor growth in *Ifnar1*$^{-/-}$ (n=7), *Xcr1*$^{cre}$*;Cxcl9*$^{fl/KO}$ (n=4), *Cxcl9*$^{fl/KO}$ (n=6), *Xcr1*$^{cre}$*;Il15ra*$^{fl/KO}$ (n=4) and *Il15ra*$^{fl/KO}$ (n=5) and Control (n=6) female mice. One experiment representative of >2 independent ones is shown. *, $p < 0.05$; **, $p < 0.01$; ***, $p < 0.001$ (unpaired *t* test).

### Supplementary Figure S5

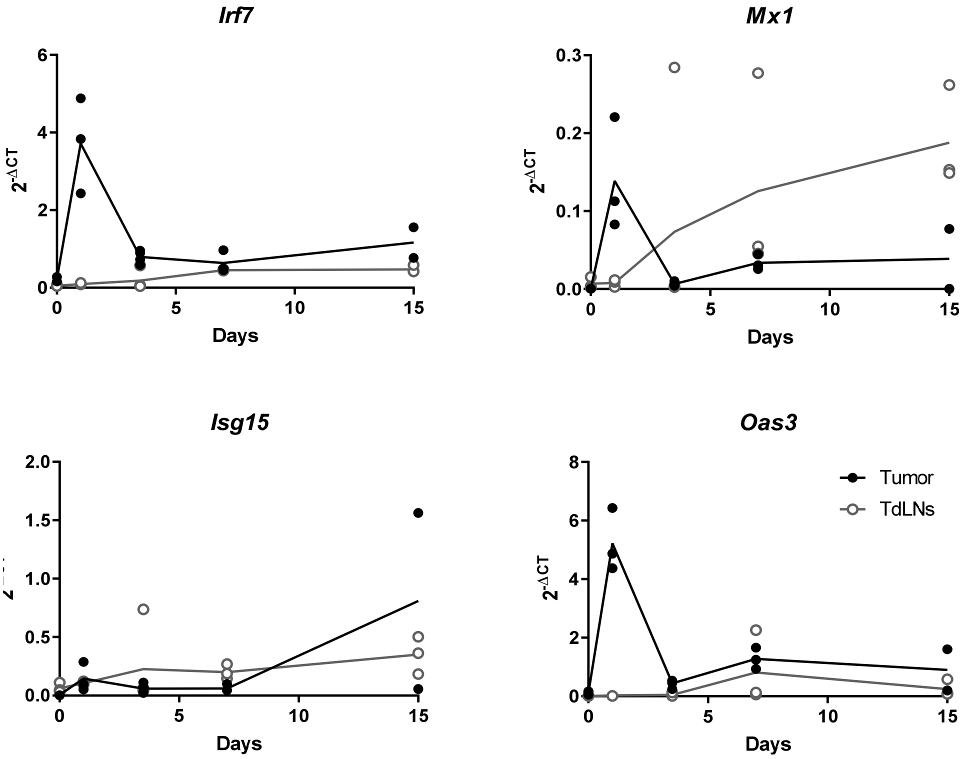


**Supplementary Figure S5.** ISGs are transcribed in tumor and TdLNs

Expression of the *Irf7*, *Mx1*, *Isg15* and *Oas3* genes in tumor and TdLNs (n=2-4) from *WT* mice.

**Supplementary Figure S6**

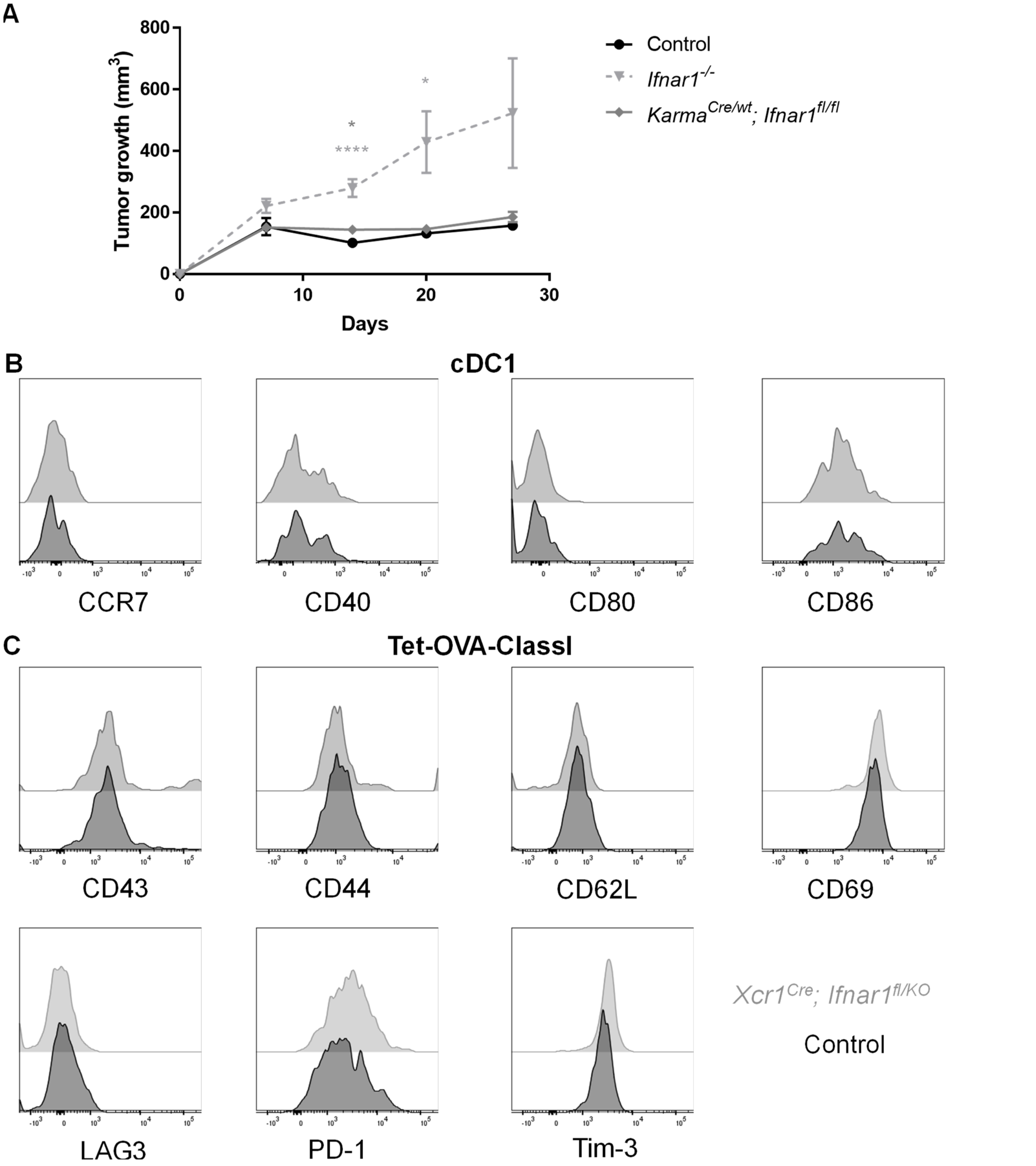


**Supplementary Figure S6.** cDC1 cell-intrinsic responses to IFN-I are dispensable for tumor control **A,** Tumor growth in *Ifnar1*$^{-/-}$ (n=8), *Karma*$^{Cre/wt}$*;Ifnar1*$^{fl/fl}$ (n=8) and control (n=7) female mice. One experiment representative of two independent ones is shown. *, P <0.05; **, P<0.01; ***, P<0.001; ****, P<0.0001 (unpaired *t* test). **B-C,** Expression of activation markers on tumor-infiltrating cDC1 (**B**) and Ag-specific CD8$^+$ T cells (**C**) at d25 post-engraftment in *Xcr1*$^{cre}$*;Ifnar1*$^{fl/KO}$ and control mice.

### Supplementary Figure S7

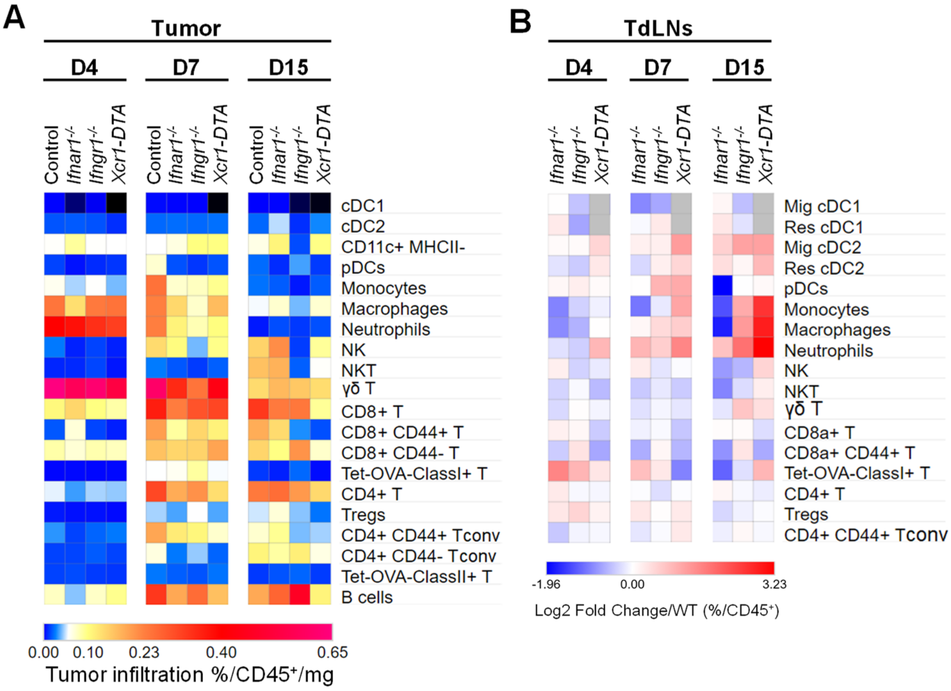


**Supplementary Figure S7.** cDC1 and IFN signaling shape the tumor immune landscape

Heatmaps representing the global immune landscapes in the tumors (**A**) and TdLNs (**B**) at d4, d7 and d15 after engraftment in *Ifnar1*$^{-/-}$, *Ifngr1*$^{-/-}$, *Xcr1-DTA* and control mice (n=3-6 mice per group). The data are shown as (mean % Population/CD45$^{+}$/mg) (A) and as Log2 Fold Changes calculated as the ratio of % Population/CD45$^{+}$ from mutant animals to WT (B). The data shown are from two independent experiments pooled together.

**Supplementary Figure S8**

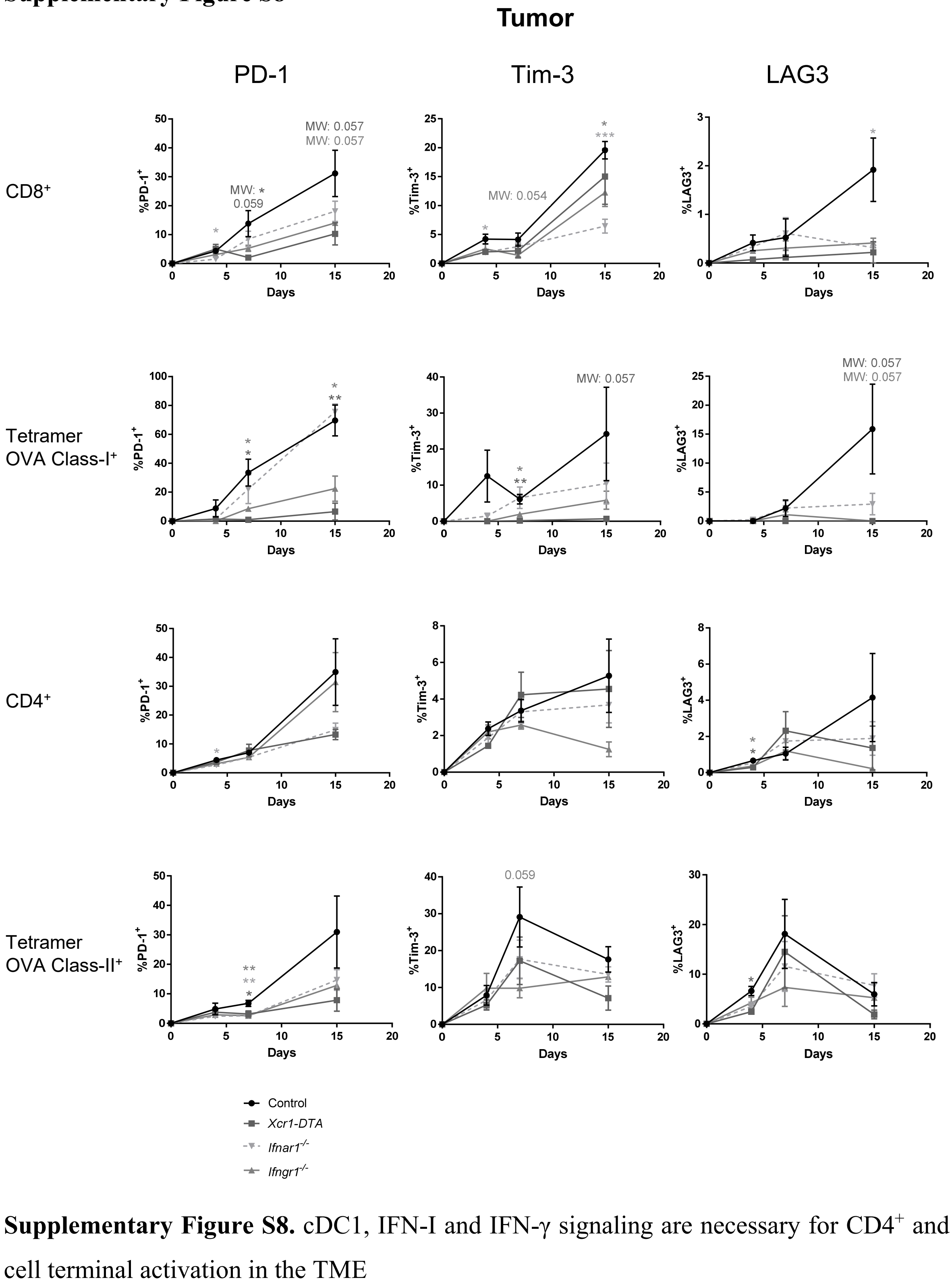


**Supplementary Figure S8.** cDC1, IFN-I and IFN-γ signaling are necessary for CD4⁺ and CD8⁺ T cell terminal activation in the TME

Frequencies of PD-1⁺; Tim-3⁺ and LAG3⁺ among CD8⁺, Ag-specific CD8⁺, CD4⁺ and Ag-specific CD4⁺ T cells at d4, d7 and d15 in the tumors from control, *Ifnar1*$^{-/-}$, *Ifngr1*$^{-/-}$ and *Xcr1-DTA* mice. The data shown (mean+/-SEM) are from two independent experiments pooled together (n=3-6 mice per group). ns, not significant (p>0.05); *, p<0.05; **, p<0.01; ***, p<0.001; (unpaired *t* test or nonparametric Mann–Whitney test [MW] when specified).

### Supplementary Figure S9

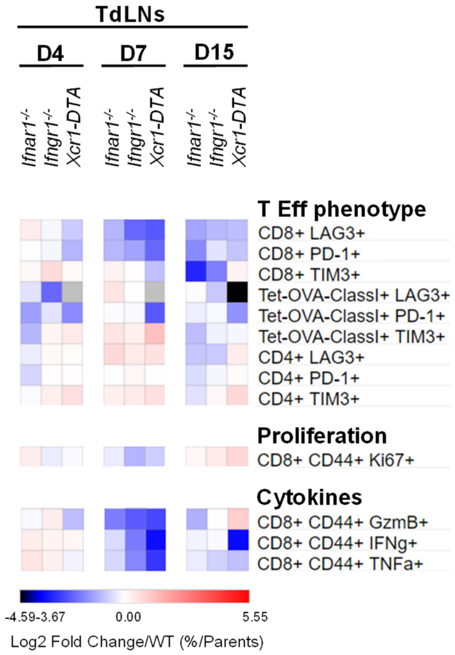


**Supplementary Figure S9.** cDC1, IFN-I and IFN-γ signaling shape TdLN immune composition, immune population activation states and effector functions

Heatmap representing effector T cell phenotype, CTL proliferation and CTL cytokine production. For cytokine production, cell suspensions were restimulated *ex vivo* with SIINFEKL peptide. The data are shown as Log2 Fold Changes of the mean %Population/Parent population of mutant animals compared to their WT counterparts (n=3-6 mice per group). The data shown are from two independent experiments pooled together.

**Supplementary Figure S10**

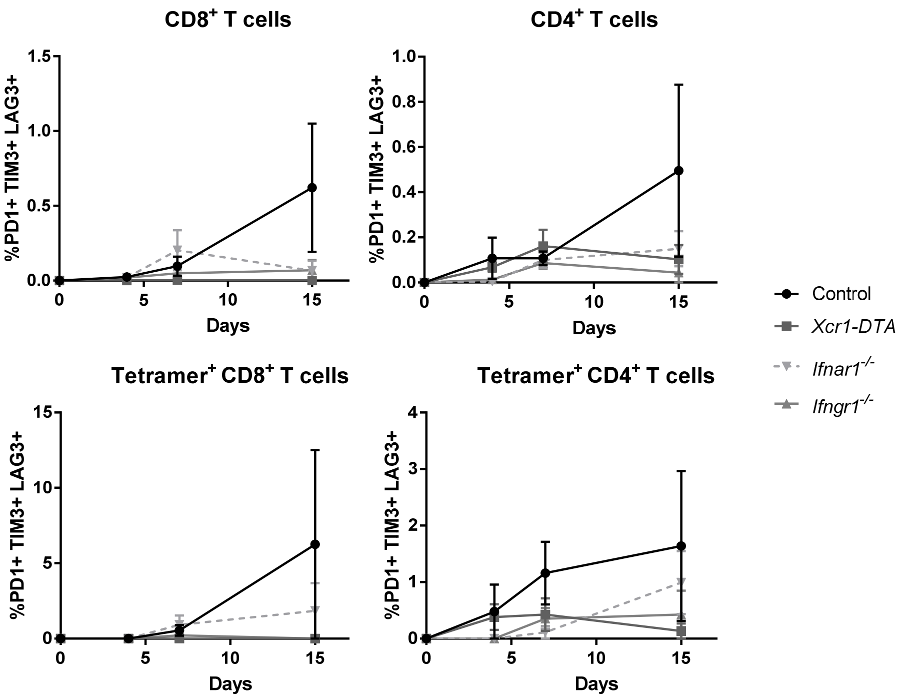


**Supplementary Figure S10.** "Exhausted-like" T cells are rare in tumors

Kinetic analysis of the percent of tumor-infiltrating cells co-expressing PD-1, Tim-3 and LAG3 among $CD8^+$, Ag-specific $CD8^+$, $CD4^+$ and Ag-specific $CD4^+$ T cells at d4, d7 and d15 in *Ifnar1*$^{-/-}$, *Ifngr1*$^{-/-}$ and *Xcr1-DTA* and in control mice. The data shown (mean+/-SEM) are from two independent experiments pooled together, each with 3-6 mice per experimental group.

**Supplementary Figure S11**

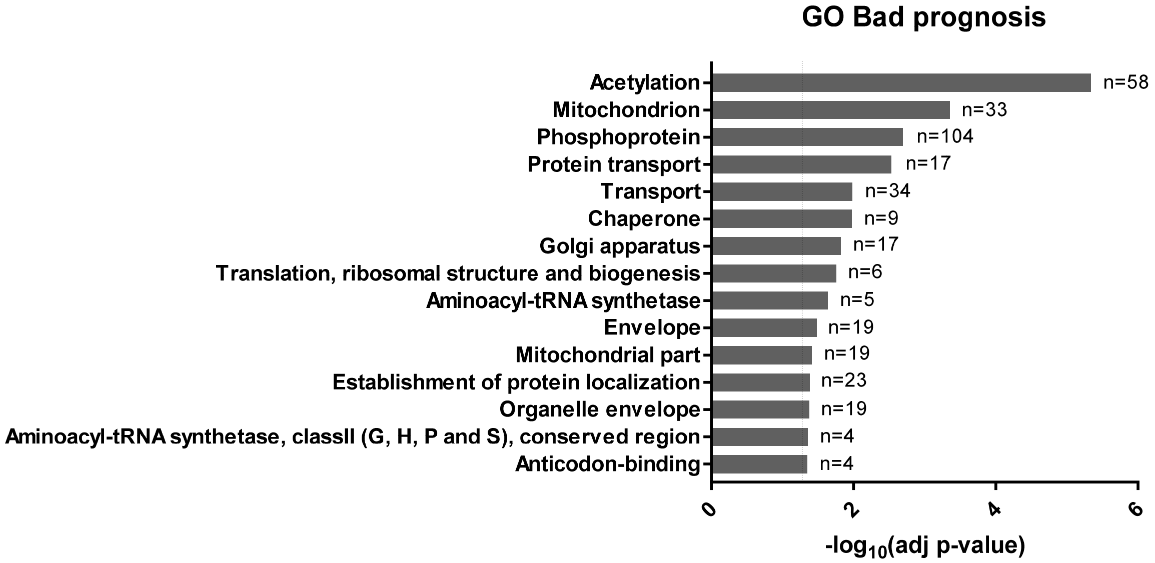


**Supplementary Figure S11.** Energy metabolism genes are associated with a bad prognosis in human breast cancer patients

Gene Ontology of the breast cancer bad prognosis gene set. The data source used and the type of analysis performed are the same as depicted in the legend of Figure 6.

**Supplementary Figure S12**

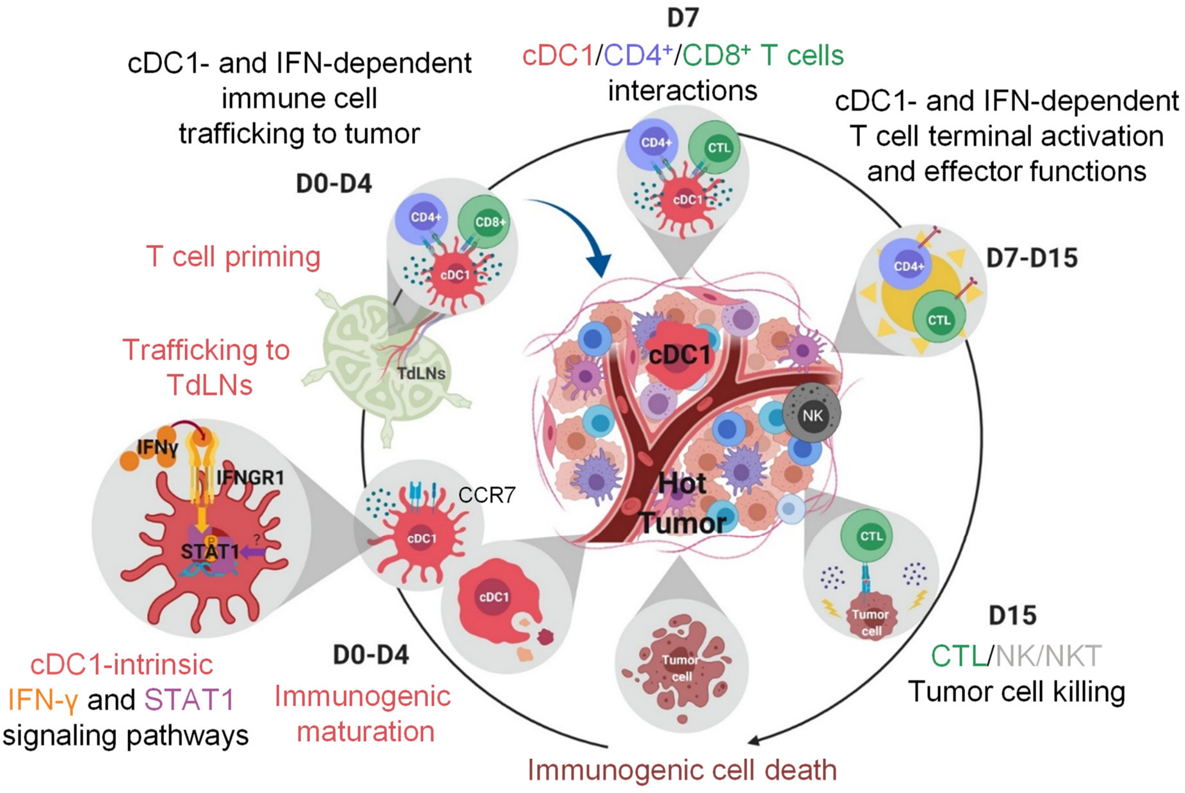


**Supplementary Figure S12.** Proposed model of cDC1 and IFN role in the immunosurveillance against breast cancer

Within the 4 first days after tumor engraftment, type I and II IFNs participate in shaping a microenvironments suitable to confer immunogenicity to cDC1 resident of tumors, or in the periphery of tumors. These cDC1 acquire CCR7 expression, which confers them their capacity to migrate into the TdLN. In the TdLN, they may prime both $CD4^+$ and $CD8^+$ T cells in type I IFN-independent manner. At day 7 in the TME, cDC1 are seen interacting simultaneously with $CD4^+$ and tumor-specific $CD8^+$ T cells, promoting their terminal differentiation and functional responses instrumental for tumor rejection. NK/NKT cells would also participate in eliminating tumor cells.
